# Clustering of RNA Polymerase II C-Terminal Domain Models upon Phosphorylation

**DOI:** 10.1101/2024.06.28.601284

**Authors:** Weththasinghage D. Amith, Vincent T. Chen, Bercem Dutagaci

## Abstract

RNA Polymerase II (Pol II) C-terminal domain (CTD) is known to have crucial roles in regulating transcription. CTD has also been highly recognized for undergoing phase separation, which is further associated with its regulatory functions. However, the molecular interactions that the CTD forms to induce clustering to drive phase separations and how the phosphorylation of CTD affects clustering are not entirely known. In this work, we studied the concentrated solutions of two heptapeptide repeats (2CTDs) models at different phosphorylation patterns, protein, and ion concentrations using all-atom molecular dynamics simulations to investigate clustering behavior and molecular interactions driving the cluster formation. Our results show that salt concentration and phosphorylation patterns play an important role in determining the clustering pattern, specifically at low protein concentrations. The balance between inter- and intra-peptide interactions and counterion coordination together impact the clustering behavior upon phosphorylation.

## INTRODUCTION

The transcription process of genetic information of DNA into RNA is mainly regulated by the RNA Polymerase II (Pol II) complex which consists of several subunits.^1, 2^ The C-terminal domain (CTD) of the largest subunit (RPB1) of Pol II is a disordered low-complexity region, that falls into the category of intrinsically disordered proteins (IDPs).^3, 4^ The CTD consists of heptapeptide (YSPTSPS) repeating units and the number of repeating units differs according to the organism.^3^ The CTD has critical roles in transcription at several steps including initiation,^5, 6^ pausing,^7, 8^ elongation,^9, 10^ splicing^11, 12^ and termination,^13, 14^ and it undergoes post-translational modifications, including phosphorylation, a key regulatory modification in these transcription steps.^3, 15^

CTD has been also recognized for its fundamental role in liquid-liquid phase separation (LLPS).^4, 16–20^ Phosphorylation of CTD impacts the LLPS formation^4, 16, 21–23^ and has been suggested as a regulator for CTD to switch between different condensates.^4, 16, 21, 23^ However, there is limited knowledge of the structural properties of CTD and the effects of phosphorylation on the structure and clustering behavior.^17, 24–26^ It has been intuitively assumed that introducing phosphorylation might extend the CTD structure due to the increased repulsion between negatively charged phosphate groups^3, 27^ and a recent study supported this prediction by identifying a more extended structure upon phosphorylation using the Small Angle X-ray Scattering (SAXS) technique.^26^ In addition, in our earlier work,^28^ we showed that phosphorylation has complex effects on the structural properties of CTD models, which were either compacted or extended depending on the CTD length and phosphorylation pattern.

The knowledge of the clustering behavior of CTD is also limited. Previous studies on crowded systems of proteins using both experimental^20, 29–32^ and computational^32–36^ methods show that protein-protein interactions play an important role in forming clusters, while the nature of such interactions for CTD clustering has not been entirely known. The clustering of CTD and formation of liquid droplets were observed by in vitro studies^4, 16^ and a recent study suggested a critical role for tyrosine interactions in cluster formation by molecular dynamics (MD) simulations and nuclear magnetic resonance (NMR) spectroscopy.^17^ How the phosphorylation affects the clustering, on the other hand, is still an open question. To obtain insights into this open question, in this work, we investigated the clustering behavior of CTD models in different phosphorylation states using all-atom MD simulations. Following our previous work,^28^ we studied the 2-heptapeptide repeating units (2CTD) model in its non-phosphorylated form, and at three different phosphorylation states. We performed simulations in three different protein concentrations to mimic a high-concentration crowded environment and three different salt concentrations to monitor the effects of ionic strength on the clustering of CTD. The initial systems were prepared using the low energy conformations of each 2CTD model that were extracted from the replica-exchange molecular dynamics (REMD)^37^ simulations performed in our recent study.^28^ After performing μs timescale MD simulations, we analyzed the clustering behavior and the molecular interactions that could drive the cluster formation. We observed that inter-peptide interactions along with ion coordination with negatively charged phosphate groups of different peptides dictate the clustering patterns of these crowded systems.

## COMPUTATIONAL METHODS

### System Preparation

For this study, we prepared multiple systems at different concentrations of 2CTD, patterns of phosphorylated serine residues, and salt concentrations. The exact models used for this study with their respective sequences, abbreviations, protein concentrations, number of peptides, and net charge per peptide are shown in Table 1. To model the concentrated systems at each protein concentration, we used initial structures from the low-energy conformations of the four 2CTD sequences that were obtained from our previous study.^28^ All the crowded systems were generated using the following protocol: For all 2CTD peptides, acetyl (ACE) and -NHCH3 (CT3) groups were utilized to cap the N-terminus and C-terminus respectively. Each 2CTD crowded system was solvated in a cubic box with each side length of the simulation box set to 100 Å. The systems were neutralized by including sodium (Na^+^) ions without additional salt (0 mM NaCl). In addition, each protein concentration was modeled at 150mM, and 300 mM NaCl concentrations to investigate the salt effect on clustering. Overall, the CHARMM-GUI server^38, 39^ was utilized to prepare and model the 36 systems with different salt concentrations, protein concentrations, and phosphorylated states as specified in Table 1.

**Table 1.**
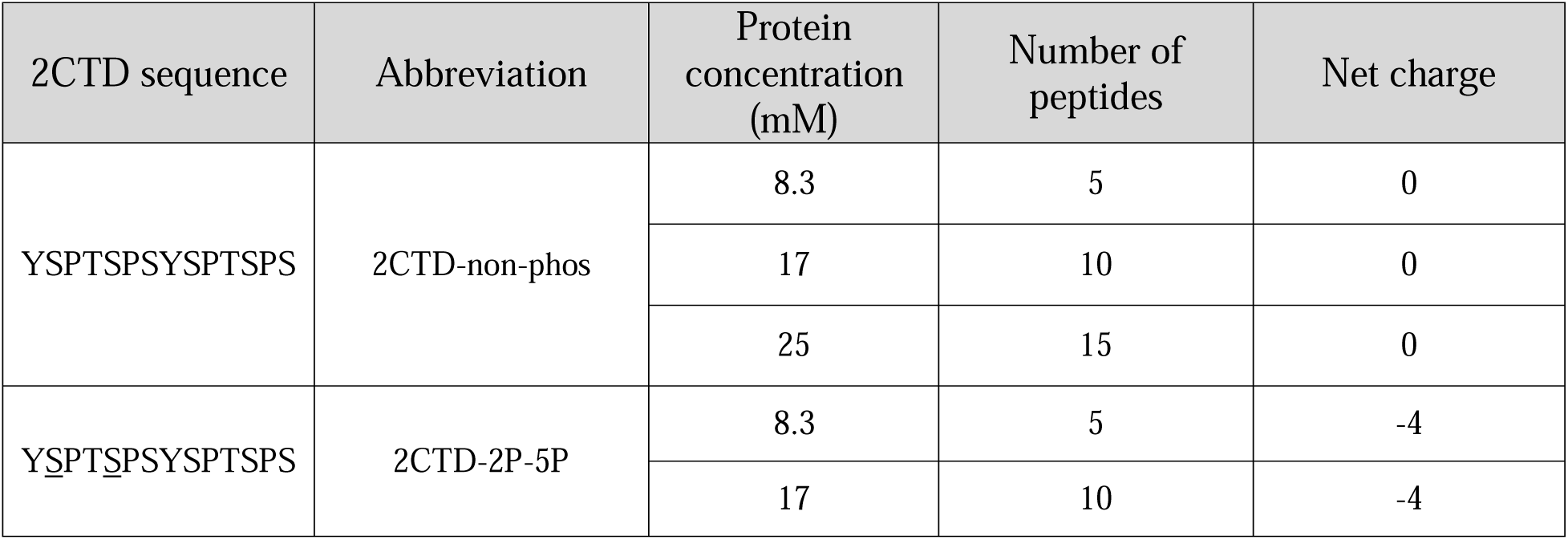
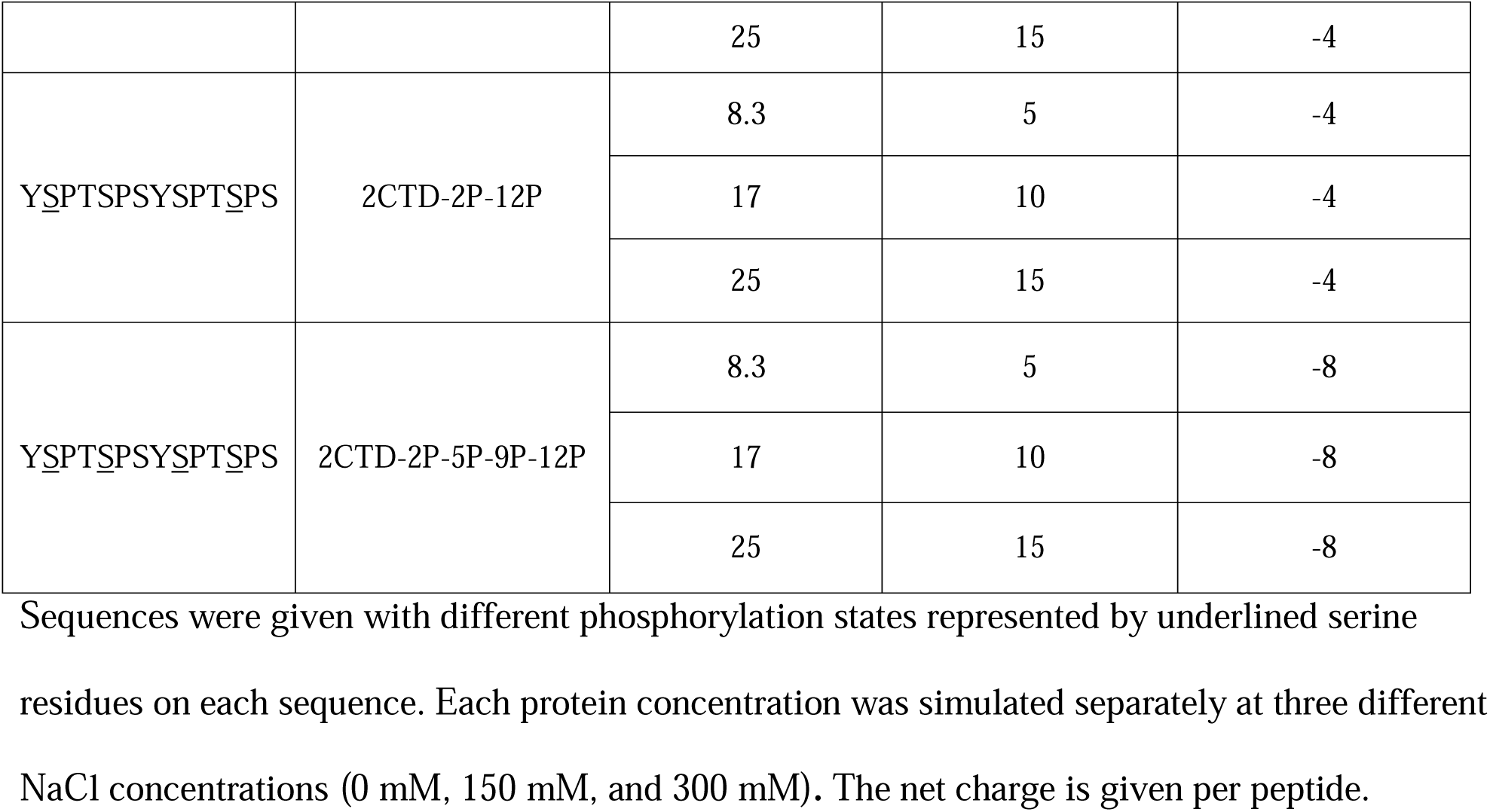
2CTD models with their sequences, abbreviations, protein concentrations, number of peptides, and net charge.

### MD Simulations

The CHARMM-modified TIP3P forcefield^40^ was utilized for the explicit water. For protein, we used CHARMM 36m^41^ force field with modifications in the nonbonded parameters between protein and water that the details were described in Huang *et al*.^41^ The energy minimization for each crowded solution was performed for 5000 steps with a tolerance of 100 kJ/mol. Then, systems were equilibrated for 1 ns while increasing the temperature from 100 to 300 K. Restraints for each 2CTD system were turned on for the initial 625 ps of the equilibration and turned off for the last 375 ps of the equilibration. During the initial 625 ps of the equilibration, both the backbone and side chains of the 2CTDs were constrained using the force constants 400 and 40 kJ/mol/nm^2^ respectively. The Lennard-Jones interactions were switched between 1.0 and 1.2 nm where the timestep was set to 1 fs for the equilibration steps.

The output configurations were extracted from the last equilibration step and utilized as the input configurations for the production runs performed for 1 µs for each system. Long-range electrostatic interactions were calculated using the particle mesh Ewald (PME)^42, 43^ algorithm with periodic boundary conditions. The Lennard-Jones interactions were switched between 1.0 and 1.2 nm. The temperature was maintained at 300 K for the production runs. The friction coefficient was set to 1 ps^-1^ for the Langevin thermostat. The time step was set to 2 fs, and frames were saved to the MD simulation trajectories at every 10 ps for the production runs. All the MD simulations were performed using OpenMM package^44^ on GPU-enhanced environment. Overall, a total of 36 µs were achieved after all the production runs.

### Analysis of MD Simulations

For the data analysis, we used the last 800 ns of the 1 µs MD trajectories from the production runs for each system. All the analyses were performed using MDAnalysis package^45^ and custom Python and C++ scripts.

We calculated the cluster size distributions using a distance cutoff value following the previous works.^33, 46^ A cluster between two peptides is counted when the distance between two alpha carbon (CA) atoms is below the cutoff of 7 Å. For example, if all 5 peptides are closer than the cutoff at 8.3 mM protein concentration at a certain frame, the cluster size outputs as 5 for that frame. The output data was used to generate the distributions of the largest cluster sizes along the simulations and percentages of each cluster size as shown in the results section.

The self-diffusion coefficients (D) were calculated from the mean squared displacements (MSD) using the Einstein formula in MDAnalysis^45^ by considering all the peptides for each 2CTD crowded system with unwrapped MD trajectories. Figure S1 in the supporting information (SI) shows how we calculated D for each crowded system by plotting MSD values against the lag times. To determine the slope of the plot, we performed a linear regression between 20 ps and 100 ps lag times. MSD values were calculated using the last 800 ns of the simulations and with a time window of 10 ps between steps which was the frequency of saving the frames during the production run and the Fast-Fourier Transform (FFT) approach^47^ implemented in MDAnalysis with the tidynamics package^48^ to improve the speed of the calculation. After the linear regression, D was calculated by dividing the slope of the fitted linear line by 2 * *dimension factor*. Here, the *dimension factor* is equal to 3, because we are considering all three coordinates of the peptides during the MSD calculation.

The radius of gyration (R_g_) for each peptide was calculated in a frame and then averaged at each frame over the number of peptides for each protein concentration. To obtain the conformational landscapes, we performed PCA on the aligned cartesian coordinates. We first aligned all the peptides in each frame along the trajectory against the initial conformation. Then, we calculated the average structure and performed a second alignment against the average structure. The cartesian coordinates of the backbone atoms of the aligned peptides were considered to perform the PCA. Only the first and second principal components (PC1 and PC2) were considered. The free energy landscapes of the PCA were obtained by applying the weighted histogram analysis method (WHAM) using the WHAM package developed by Grossfield lab.^49^ For the PCA, MATLAB^50^ (along with PC1 and PC2, which served as the reaction coordinates of the 2CTD systems) was utilized to plot the free energy landscapes. To extract the frames with the minimum energy conformations, first, for each peptide within each system we identified the lowest energy conformation located at the energy minimum of the corresponding PCA plot. Then, we selected the frame corresponding to the minimum energy conformations of each peptide. Therefore, we selected five frames, each with the minimum energy conformation of one of the five peptides in the 8.3 mM concentrated CTD systems. Then, the frame with the highest number of low-energy conformations among the five selected frames was extracted for each system. We used the last 800 ns of simulations for the PCA and the last 100 ns for the extraction of the low-energy conformations.

Contact maps were generated by considering the inter-peptide contacts below or equal to 5 Å for 2CTD systems at 8.3 mM protein and 150 mM NaCl concentrations. Contacts were averaged for each residue pair over the number of the peptide pairs and number of the frames of the trajectories.

### Data Availability

We provided the scripts for the analyses of the simulations and the snapshots with the lowest energy conformations for each simulation at 8.3 mM protein concentrations in a GitHub repository (https://github.com/bercemd/ctd_clustering).

## RESULTS

In this study, we investigated the clustering patterns of the non-phosphorylated 2CTD model and 2CTD models at different phosphorylation states. We obtained insights into the intermolecular interactions and effects of phosphorylation within heptapeptide repeats towards developing an understanding of the LLPS occurring in full human CTD. In the main text, we will only show the results related to the lowest protein concentration of 2CTD crowded systems which is 8.3 mM. All the results related to the other two high protein concentrations (17 mM and 25 mM) are shown in the SI.

### 2CTD systems showed varied degrees of clustering depending on the phosphorylation pattern

Figure 1 shows the largest cluster size distributions (1a-c) and the cluster size percentages (1d-f) at 8.3 mM protein concentration and three different NaCl concentrations. For 2CTD-non-phos, the largest cluster size distributions did not show any significant changes (average stays around 2) across different ion concentrations (Figure 1a-c), and higher percentages were observed for monomers and dimers (Figure 1d-f).

**Figure 1.**
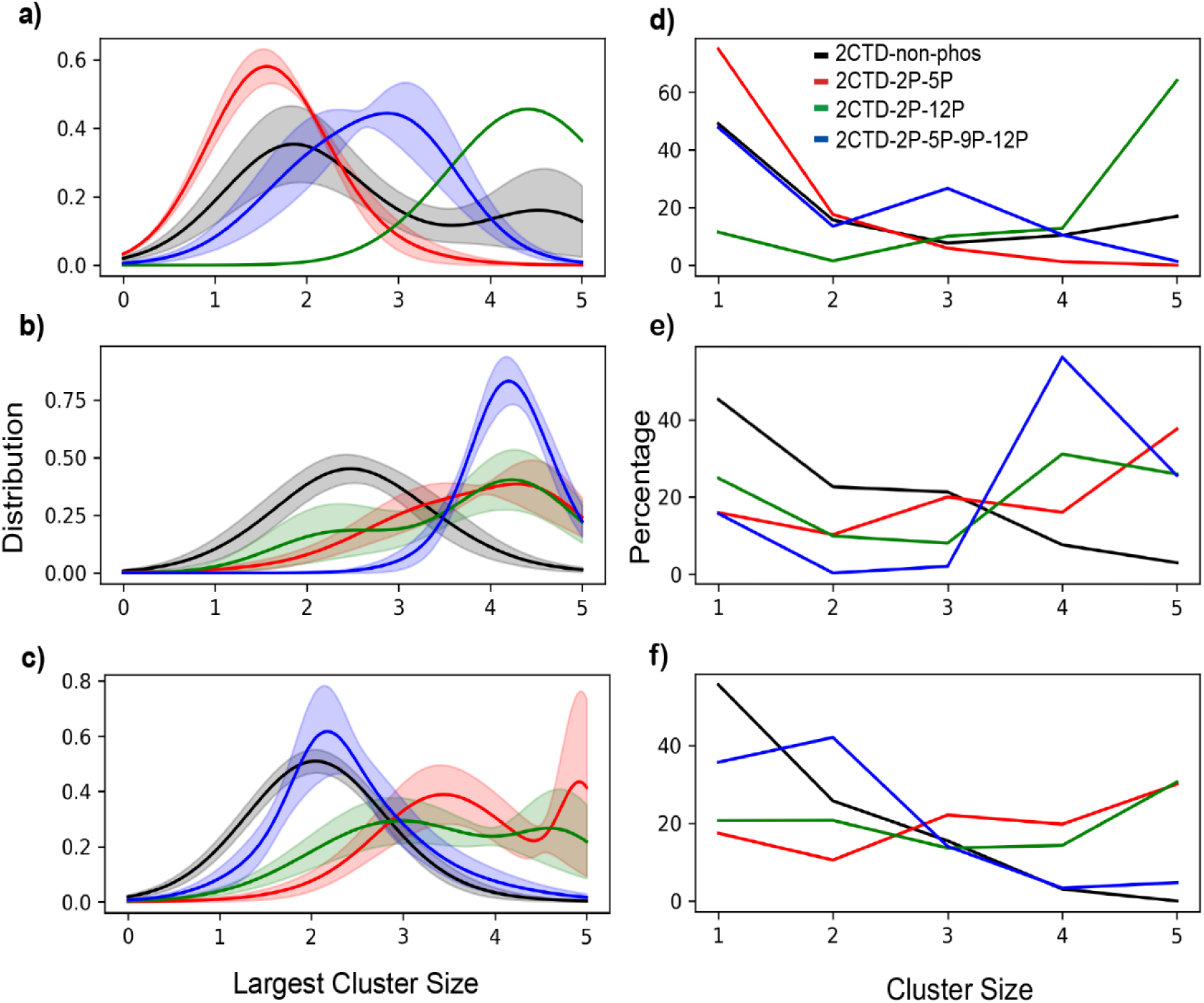
Cluster distribution and cluster size percentages in 2CTD crowded systems at 8.3 mM protein concentration. The largest cluster size distributions are shown at a) 0, b) 150, and c) 300 mM NaCl. Percentages of each cluster size are shown at d) 0, e) 150, and f) 300 mM NaCl. Standard errors [for panels (a), (b) and (c)] were calculated by splitting the last 800 ns of the 1 µs trajectory into 160 ns small trajectories for each system. For 2CTD-2P-12P in panel (a) the standard error is not shown because approximately for the last 320 ns the largest cluster size remained constant at 5. The colors of the curves are the same as in panel (d) for all the other panels.

Phosphorylated 2CTD systems show more diverse trends of the largest cluster size distributions at each ion concentration. Generally, phosphorylated systems display higher cluster sizes compared to the non-phosphorylated state except for the case of 2CTD-2P-5P at 0 mM NaCl concentration in Figures 1a and d. Both 2CTD-2P-5P-9P-12P and 2CTD-2P-5P have cluster sizes increased by increased salt concentration from 0 to 150 mM, then reduced by increasing the salt concentration even more to 300 mM for 2CTD-2P-5P-9P-12P and stayed around the same for 2CTD-2P-5P. This suggested that Na^+^ ions may form bridging electrostatic interactions with phosphate groups, which increase the clustering at 150 mM concentration, while in a higher concentration of NaCl electrostatic interactions are reduced due to the charge screening. For 2CTD-2P-12P, on the other hand, we observed the largest clusters at the lowest ion concentration suggesting that electrostatic interactions at higher ion concentrations are hindered potentially because of a conformational change. In addition, diffusion coefficients of peptides correlate with the observed clustering showing that increased clustering reduced translational diffusion of peptides (Table 2). The clustering analysis supported by diffusion coefficient calculations, overall suggests that different phosphorylated systems have complex patterns of clustering at different salt concentrations.

**Table 2.** Self-diffusion coefficients of 8.3 mM 2CTDs at different salt concentrations.

| NaCl<br>concentration<br>(mM) | Diffusion Coefficients ($10^{-2} \text{ \AA}^2 \text{ ps}^{-1}$) | | | |
| --- | --- | --- | --- | --- |
|  | 2CTD-non-phos | 2CTD-2P-5P | 2CTD-2P-12P | 2CTD-2P-5P-9P-12P |
| 0 | 3.81 | 3.39 | 2.01 | 2.48 |
| 150 | 3.83 | 2.34 | 2.30 | 1.67 |
| 300 | 3.92 | 2.23 | 2.17 | 2.16 |

At higher protein concentrations (at 17 and 25 mM), we observed a larger degree of clustering, especially at higher NaCl concentrations (at 150 and 300 mM). Figures S2 and S3 show the largest cluster size distributions and cluster size percentages for the 2CTD systems at 17 and 25 mM protein concentrations, respectively. For both concentrations, we observed higher values for the largest cluster sizes for both non-phosphorylated and phosphorylated crowded systems compared to 8.3 mM protein concentration. All the peptides tend to form one large cluster, regardless of phosphorylation states at high ion concentrations. These observations suggest that 17 and 25 mM protein concentrations might be unrealistically large and potentially cause aggreagation as we also observed in substantial decrease in diffusion coefficients especially for 25 mM concentration (Tables S1-2). Therefore, we focus on the lowest protein concentration (8.3 mM) and discuss the specific characteristics of that concentration in the following sub-sections.

### Crowding and ion concentration alter the conformations and impact clustering

We observed that phosphorylation and ion concentration have complex effects in clustering as one phosphorylated system has the largest cluster sizes at 0 mM salt concentration while the other two show an opposite trend from 0 to 150 mM salt concentrations. To understand these clustering patterns, we need to investigate the structural properties of these 2CTD crowded systems. For that purpose, we analyzed the conformational changes for peptides in each 2CTD system.

In order to investigate the conformational space of 2CTD systems, we first generated the average R_g_ distributions over the number of peptides for each 2CTD system at three different NaCl concentrations shown in Figure 2. At 0 mM salt concentration (only in the presence of neutralizing counterions), we observed that 2CTD-2P-5P and 2CTD-2P-12P extended their structures while 2CTD-2P-5P-9P-12P contracted compared to 2CTD-non-phos in Figure 2a. This observation deviates for 2CTD-2P-12P system from the results in our earlier work^28^, in which we observed contraction for 2CTD-2P-12P compared to 2CTD-non-phos using REMD simulations of isolated single peptides. Expansion of 2CTD-2P-12P in the concentrated system compared to the isolated system suggests that crowding affects the conformation in a way that inter-peptide interactions dominate over the interactions with nearby residues (intra-peptide). This conformational change consequently resulted in increased clustering observed in Figures 1a and d. At higher salt concentrations, 2CTD-2P-12P showed more contracted conformations (Figure 2b-c) suggesting that not only crowding but also salt concentration altered the conformation of this peptide. Average R_g_ distributions for 17 and 25 mM protein concentrations are shown in Figure S4 and S5 respectively at three different salt concentrations. The average R_g_ distribution patterns were similar to the patterns of 8.3 mM protein concentration (except at 0 mM NaCl concentration) at all three salt concentrations for all the crowded systems.

**Figure 2.**
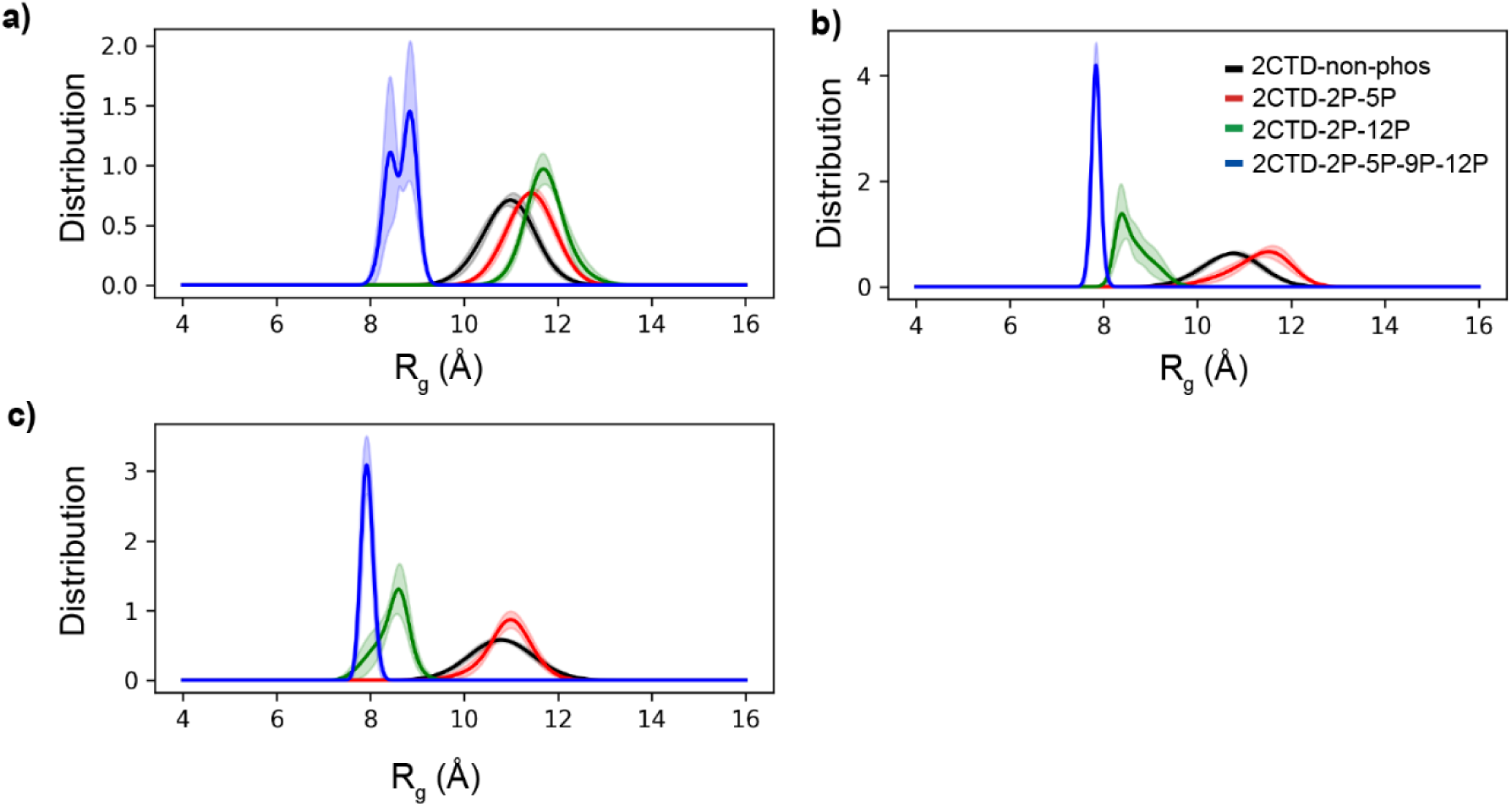
Average radius of gyration (R_g_) density distributions for 2CTD crowded systems at 8.3 mM protein and (a) 0, (b) 150, and (c) 300 mM NaCl concentrations. The colors of the curves are the same as in panel (b) for all the other panels. Standard errors were calculated by splitting the last 800 ns of the 1 µs trajectory into 160 ns small trajectories for each system.

To obtain a deeper understanding of the conformational space of 2CTD-2P-12P compared to other three 2CTD crowded systems, we generated free energy landscapes of PCA using the cartesian coordinates of backbone atoms of peptides as described in the Methods section and extracted the frames with the low energy conformations shown in Figure 3 for 2CTD-2P-12P, Figure S6, S7 and S8 in the SI for 2CTD-non-phos, 2CTD-2P-5P-9P-12P and 2CTD-2P-5P respectively at three NaCl concentrations. Figure 3a shows the extended peptides for the 2CTD-2P-12P crowded system at 0 mM NaCl concentration which confirmed the observed R_g_ behavior in Figure 2a. Additionally, we saw that 2CTD-2P-12P peptides are contracted at 150 mM and 300 mM NaCl concentrations in Figure 3b and c respectively, compared to at 0 mM NaCl (Figure 3a) which also aligned with the R_g_ behavior of 2CTD-2P-12P in Figure 2b and c. For the other three 2CTD systems, we confirmed the extended (for 2CTD-non-phos and 2CTD-2P-5P) and contracted (for 2CTD-2P-5P-9P-12P) peptide conformations at three NaCl concentrations in Figures S6-S8.

**Figure 3.**
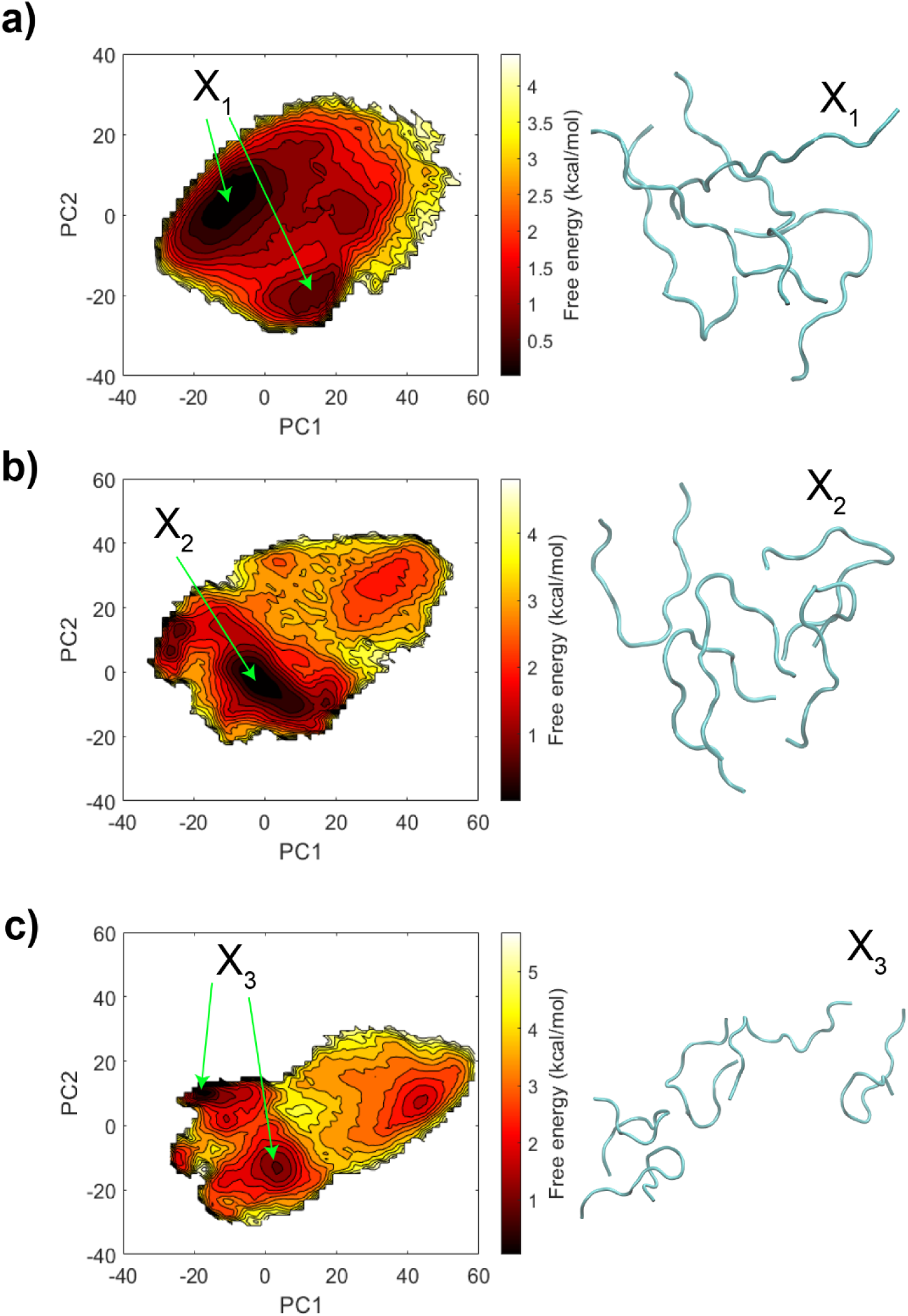
Free energy landscapes using PC1 and PC2 as reaction coordinates from the PCA using cartesian coordinates for 2CTD-2P-12P crowded models at 8.3 mM protein and (a) 0, (b) 150, and (c) 300 mM NaCl concentrations. In addition, X_1_-X_3_ represent the frames that have low energy conformations at each salt concentration. X_1_ and X_3_ have conformations at two energy minima.

### Hydrogen bonding and counterion interactions drive the clustering of 2CTD systems

In order to determine which interactions drive cluster formation in the 2CTD crowded systems, we first calculated the total number of H-bonds associated with peptides including both intra-peptide and inter-peptide H-bonds. Figure 4 represents the distributions of the total number of H-bonds associated with peptides at 0, 150 and 300 mM NaCl concentrations.

**Figure 4.**
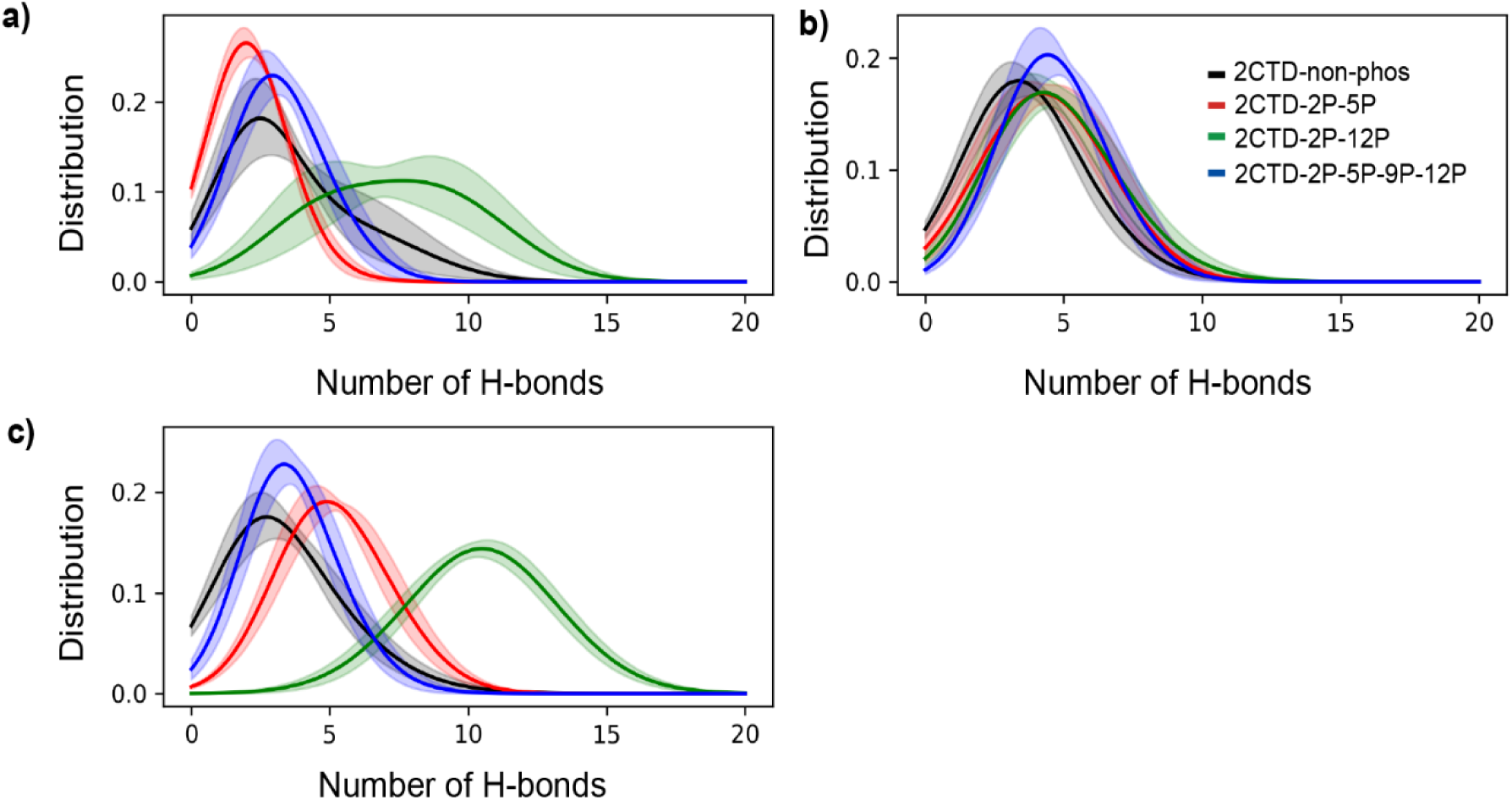
Distributions of the total number of H-bonds associated with peptides (both intra and inter) for 2CTD systems at 8.3 mM protein and (a) 0, (b) 150, and (c) 300 mM NaCl concentrations. The colors of the curves are the same as in panel (b) for all the other panels. Standard errors were calculated by splitting the last 800 ns of the 1 µs trajectory into 160 ns small trajectories for each system.

When we introduced the phosphorylation, the total number of H-bonds associated with peptides increased at all NaCl concentrations compared to 2CTD-non-phos, except for 2CTD-2P-5P at 0 mM NaCl concentration (Figure 4a). This might be the reason why 2CTD-2P-5P showed lower cluster sizes compared to 2CTD-non-phos at 0 mM NaCl concentration in Figure 1a. In addition, we observed a higher total number of H-bonds associated with peptides for 2CTD-2P-12P compared to the other three 2CTD crowded systems at 0 mM and 300 mM NaCl concentrations in Figure 4a and c respectively. Furthermore, we observed a higher number of H-bonds at 300 mM NaCl compared to 0 mM, which is not consistent with our earlier observation of a higher largest cluster size for 2CTD-2P-12P at 0 mM NaCl compared to 300 mM in Figure 1a. In order to understand the contribution of H-bonds to the clustering, we need to separate the total number of H-bonds associated with peptides into intra-peptide and inter-peptide H-bonds as shown in Figure 5 for 2CTD-2P-12P. We observed a higher number of inter-peptide H bonds (Figure 5c) compared to intra-peptide H-bonds (Figure 5b) for 2CTD-2P-12P at 0 mM NaCl concentration compared to 300 mM, which validated the higher largest cluster size at 0 mM NaCl concentration. The extended peptides (Figure 3a) and two near-terminal phosphorylated serine residues caused the formation of more inter-peptide H-bonds at 0 mM NaCl concentration. On the other hand, the higher number of intra-peptide H-bonds for 2CTD-2P-12P at 300 mM NaCl concentration (see Figure 5b) was expected as this peptide is more contracted at 300 mM NaCl in Figure 3c.

**Figure 5.**
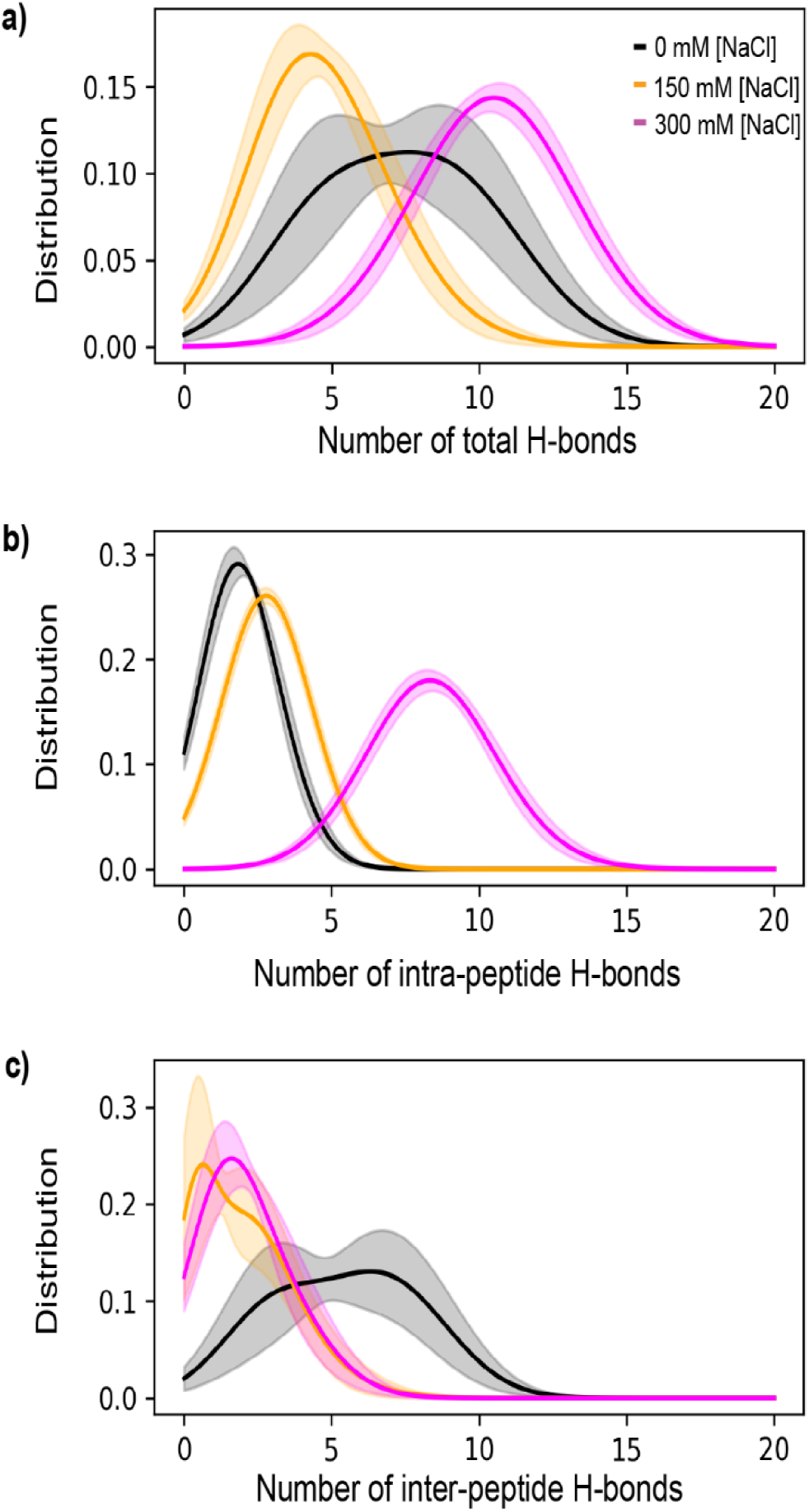
Distributions of the number of (a) total, (b) intra-peptide, and (c) inter-peptide H-bonds for 2CTD-2P-12P systems at 8.3 mM protein and different salt concentrations. The colors of the curves are the same as in panel (a) for all the other panels. Standard errors were calculated by splitting the last 800 ns of the 1 µs trajectory into 160 ns small trajectories for each system.

For the other three 2CTD crowded systems, the separation of intra-peptide and inter-peptide H-bonds is shown in Figure S9-S11 in the SI. We observed a correlation between inter-peptide H-bonds and clustering even for the contracted peptide conformations. For example, we saw a higher largest cluster size distribution for the 2CTD-2P-5P-9P-12P system, in which the peptides are contracted (Figure S7b), at 150 mM NaCl concentration in Figure 1b and this is in agreement with an increment in the number of inter-peptide H-bonds distribution in Figure S10c and clustering of peptides represents by the low energy conformations in Figure S7b. Moreover, the increments of the number of inter-peptide H-bonds for 2CTD-2P-5P at 150 and 300 mM NaCl concentrations compared to 0 mM NaCl concentration in Figure S11c, agree with the largest cluster size distributions in Figure 1b-c respectively.

A recent study by Flores-Solis, D. *et al*.^17^ which was based on both two-dimensional NMR experiments and MD simulations showed that tyrosine-proline (Y-P) interactions play an important role in driving the LLPS in human Pol II CTD and other low-complexity proteins at physiological salt concentration without phosphorylation. On that note, we calculated contact maps to obtain a perspective of residue-residue interactions which contributed to the clustering. Figure 6 represents the contact maps for the four 2CTD crowded systems which include only the inter-peptide contacts at the physiological salt concentration (150 mM). For 2CTD-2P-5P-9P-12P, 2CTD-2P-12P, and 2CTD-2P-5P, we observed a large number of inter-peptide contacts between residues specifically around phosphorylated serine residues in Figure 6b-d respectively. This confirms that even with contracted peptide conformations, crowded systems can form inter-peptide contacts to generate large cluster sizes. In addition to that, we observed additional enrichment of tyrosine interactions (including Y-P interactions) for the 2CTD-non-phos system rescaled in Figure S12.

**Figure 6.**
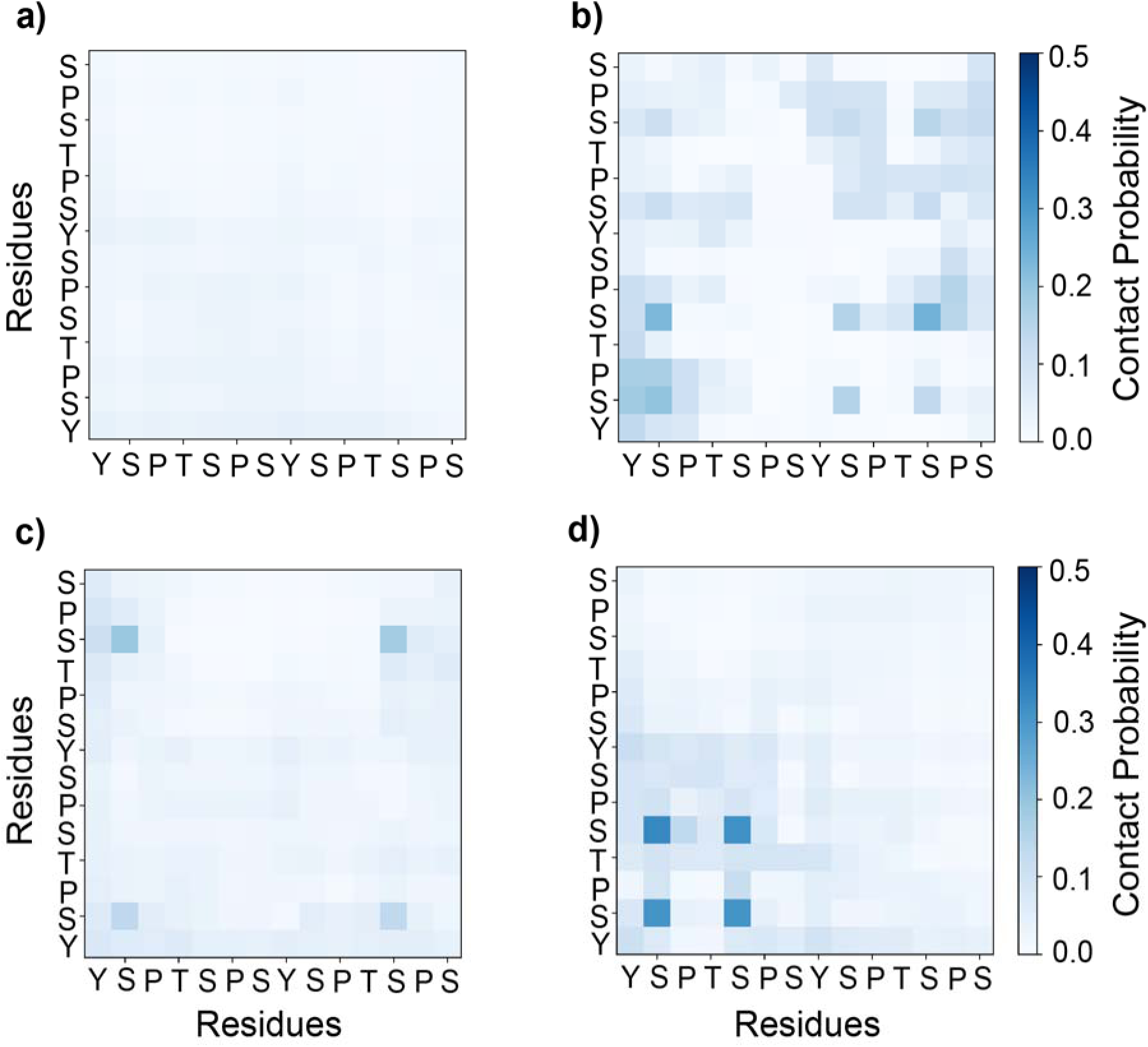
Contact maps for inter-peptide contacts between residues within 5 Å at 8.3 mM protein and 150 mM NaCl concentrations for (a) 2CTD-non-phos, (b) 2CTD-2P-5P-9P-12P, (c) 2CTD-2P-12P, and (d) 2CTD-2P-5P systems. Contacts were averaged for each residue pair over the trajectory and the number of peptide pairs.

To understand how electrostatic interactions play a role in the clustering of 2CTD peptides, we focus on phosphate groups and their interactions with Na^+^ ions. In our earlier study,^28^ we observed that Na^+^ interacts with negatively charged oxygens forming a bridge between multiple phosphate groups. To quantify Na^+^ ion bridging for the phosphate groups, we generated the distributions of distances between inter-peptide P atoms for the three phosphorylated 2CTD systems at the physiological salt concentration. Figure 7a shows that, for all three phosphorylated 2CTD systems, there is a peak around 5 Å which provides an ideal distance for Na^+^ to form coordination between the phosphate groups of two different peptides. Figure 7b shows a snapshot with low energy conformations according to the PCA plot in Figure S7b for the 2CTD-2P-5P-9P-12P system. The Na^+^ ions are in close distance with the phosphate groups from two segments (peptides) that are within around 5 Å distance confirming the Na^+^ ion bridging of the inter-peptide phosphates. These electrostatic interactions help to form inter-peptide contacts and stabilize the large clusters in phosphorylated 2CTD systems at physiological salt concentration.

**Figure 7.**
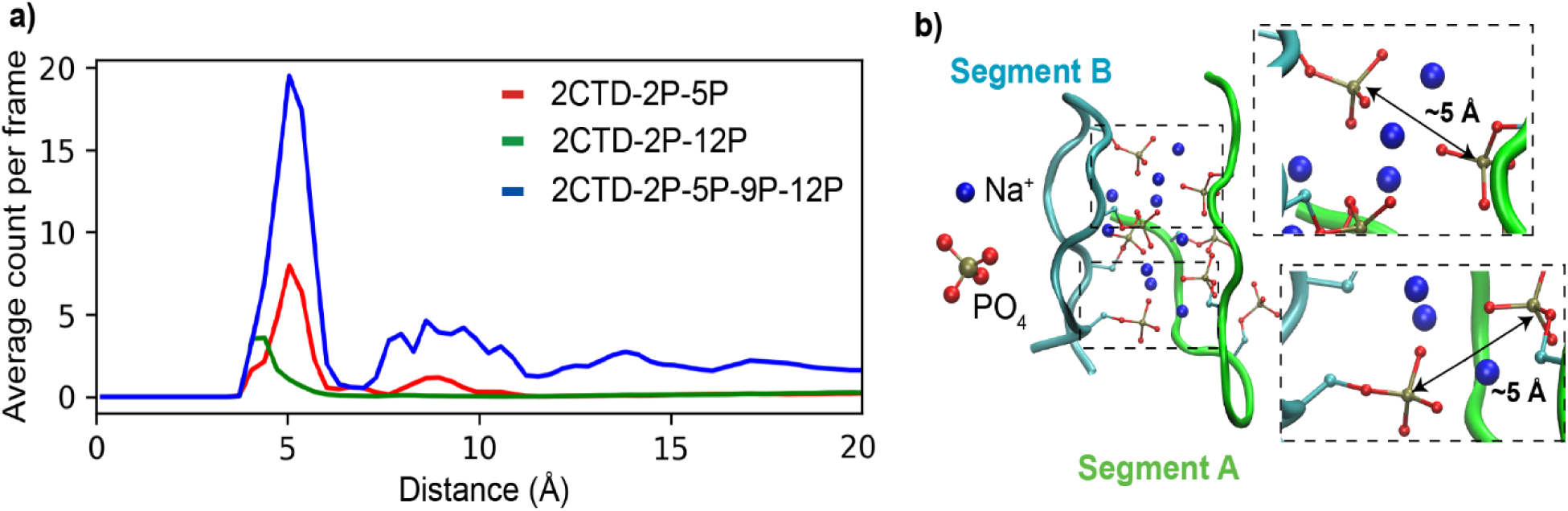
a) Distributions for distances between inter-peptide P atoms of phosphorylated 2CTDs at 8.3 mM protein and 150 mM NaCl concentrations. The average count per frame for each system was calculated by counting the number of occasions of two different peptides occupying the distance between P atoms at each specific distance bin and dividing it by the total number of frames, and b) a snapshot of 2CTD-2P-5P-9P-12P system at 8.3 mM protein and 150 mM NaCl concentrations with low energy conformations representing the distances between two P atoms of two different peptides (segment A and segment B) allowing the phosphate groups to coordinate Na^+^ ions to form clusters. Only Na^+^ ions between segments within 3 Å of both segments are shown.

### Overall effects of salt concentration and phosphorylation on clustering

In general, for higher protein concentrations (17 mM and 25 mM), large clusters formed regardless of the phosphorylation state, especially at high salt concentrations with minimal deviations within the cluster distributions (Figures S2-S3). However, there are complex effects on clustering from phosphorylation and salt concentration specifically at the lower protein concentration. Figure 8a displays a heatmap representation of how the phosphorylation and salt concentration affect the average largest cluster size for the 2CTD systems at 8.3 mM protein concentration. The average largest cluster size for 2CTD-non-phos was around below 3 at every NaCl concentration. Moreover, all three phosphorylated 2CTD systems formed higher average largest cluster sizes compared to the non-phosphorylated state at the physiological salt concentration (150 mM). Additionally, the clustering was decreased or unchanged with increasing salt concentration from 150 mM to 300 mM, suggesting that charge screening impacts the clustering behavior.

**Figure 8.**
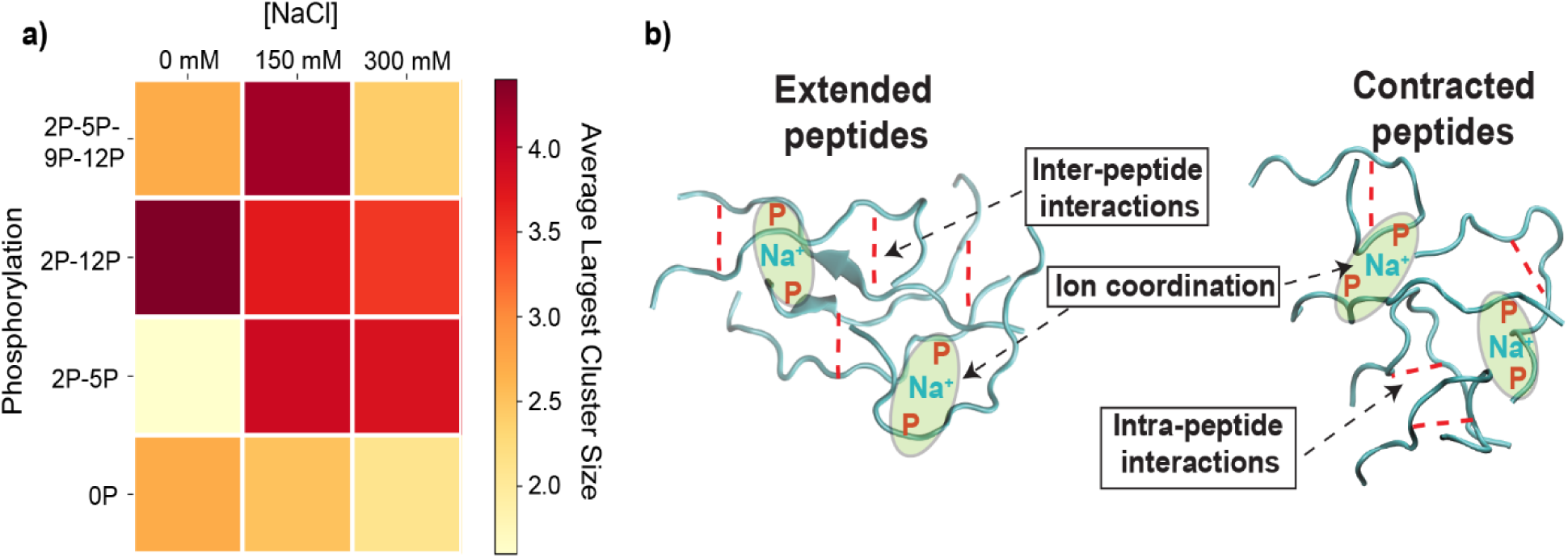
a) Heatmap for the overall effect of phosphorylation level and NaCl concentration on the clustering of 2CTD systems at 8.3 mM protein concentration and b) General model representation of how the inter-peptide interactions and ion coordination induce the formation of clusters with extended and contracted peptides.

Figure 8b shows the general model which represents how the inter-peptide interactions and ion coodinations induce the formation of the clusters. Extended peptides form clusters using both inter-peptide interactions and ion coordination while contracted peptides form clusters using mainly ion coordination upon phosphorylation. Ion coordination induces inter-peptide interactions in contracted peptides as confirmed by the contact maps in Figure 6. In summary, the clustering of 2CTD peptides depends on the phosphorylation patterns that support a combination of inter-peptide H-bondings and electrostatic interactions formed via counterion bridging.

## DISCUSSION

CTD of Pol II has been suggested to form clusters that lead to phase separation.^4, 16, 17^ In this study, we investigated the clustering behavior of CTD models using all-atom MD simulations. The CTD models used in this study consisted of two heptapeptide repeats as simple models to provide insights into the intermolecular interactions that would potentially cause clustering in longer CTDs in humans or yeast. For the non-phosphorylated CTD, we observed cluster formation in a lesser amount than the phosphorylated CTD models. The most prevalent intermolecular interactions were observed with the tyrosine residues in the heptapeptide repeats when serine residues are not phosphorylated. Since the intermolecular H-bond contribution is not substantial (Figure S9), hydrophobic interactions between the tyrosine ring and other residues may be the main contributor to the cluster formation observed for the 2CTD-non-phos. The study by Flores-Solis *et al.* on CTD clustering by both experiments and simulations suggested that tyrosine interactions are abundant in CTD clustering.^17^ They especially focused on Tyr-Pro interactions that are also observed by MD simulations in their study. Tyrosine interactions were also observed between CTD and the mediator complex that forms condensates with Pol II.^51^ The predominance of the tyrosine interactions in the 2CTD-non-phos from our simulations is aligned with the literature that reports an abundance of tyrosine interactions in CTD clustering.

Non-phosphorylated CTDs were shown to form liquid droplets in vitro and phosphorylation caused the dispersion of these droplets.^4^ Phosphorylated CTDs were also recruited to the droplets formed by other proteins, especially observed by kinases of positive transcription factor b (P-TEFb).^16^ Such interchanges between different condensates as a function of phosphorylation level have been proposed as a mechanism by which post-translational modifications regulate the transition from initiation to the elongation stage.^21^ However, the molecular interactions that change the clustering behavior of CTD upon phosphorylation are understudied. In our study, MD simulation results suggest that phosphorylation causes the formation of clusters via electrostatic interactions with Na^+^ ions that bridge intermolecular phosphate groups. These newly formed electrostatic interactions increase the number of clusters compared to the 2CTD-non-phos system. Earlier studies suggested an interchange of the interactions from hydrophobic to electrostatics in nature upon phosphorylation of CTD.^4, 16^ However, these studies indicated the formation of electrostatic interactions between negatively charged phosphorylated CTD and other positively charged IDPs, without mentioning any electrostatic interactions mediated by ions. Our observation of Na^+^ ion bridging could be related to the size of the CTD that a human or yeast CTD with higher numbers of heptapeptide repeats may behave differently compared to our 2CTD model with two heptapeptide repeats. On the other hand, we also note that ion-mediated condensations were observed for highly charged nucleic acids in the presence of Mg^2+^ ion more prevalently,^52–54^ but also observed for Na^+^ ions in high concentration in crowded systems.^52, 54^ Therefore, the role of counterion interactions in clustering and condensate formation by phosphorylated CTDs might be underestimated and needs to be considered by future studies.

The ion-mediated interactions observed in our previous study^28^ caused more compact structures for phosphorylated CTDs through bending to coordinate the counterion by multiple phosphate groups. Other computational works on IDPs showed similar observations that counterions mediate the inter- and intra-molecular interactions.^55, 56^ All these studies were conducted using AMBER or CHARMM force fields, which have been systematically parameterized and extensively validated against experimental observations^41, 57–59^ suggesting their strong ability to provide accurate results for intermolecular interactions. However, in a highly charged concentrated system, the static charges in the standard force fields may have limitations in describing the electrostatics upon the polarization of charged atoms induced by the environment. To account for charge redistribution due to the highly charged concentrated systems and to increase the accuracy of the results, polarizable force fields^60, 61^ could be used instead of standard force fields with fixed charges. Additionally, our observations of counterion-mediated clustering could be indirectly validated using experimental methods like NMR or FRET. Increased clustering upon phosphorylation may decrease the NMR relaxation times or increase the FRET efficiencies, and the ion dependency could be monitored by modifying the ion concentration.

We performed this study to understand basic interaction networks that cause clustering of CTD models at different phosphorylation levels. However, we note that our systems have two heptapeptide repeats, which are much shorter than the CTDs observed in humans or yeast, which have 52 and 26 repeats, respectively. Studies suggest that phase separation occurs when CTD has around more than 10 repeats, which is still far longer than our model systems.^4, 22^ Therefore, although we predicted that 2CTD systems form clusters, to study phase separation we need to work with longer CTD systems. This causes a limitation in studying phase separation computationally at the atomistic detail as the system sizes increase substantially with the increased number of repeats. One way to circumvent this limitation is to switch to coarse-grained representation as it was routinely done to study the phase separation of IDPs.^62–65^

## CONCLUSIONS

In this computational study, we report the cluster formation of 2CTD models upon phosphorylation at different salt concentrations. This study helped us to obtain a fundamental understanding of the intermolecular interactions that are potentially linked to LLPS formation of Pol II CTD which was reported previously in the literature. Phosphorylation caused complex effects on the clustering specifically at low protein concentration depending on the salt concentration. For phosphorylated CTDs, clusters are either formed with contracted peptides or extended peptides by forming inter-peptide interactions (H-bonds) and coordination with Na^+^ ions in between phosphate groups of different peptides. At higher salt concentrations, charge screening occurs especially for the multiple phosphorylated CTD systems. Future directions will focus on developing and applying coarse-grained models to study phase separation of full human RNA Pol II CTD in a computationally accessible timescale.

## ASSOCIATED CONTENT

### Supporting Information

The Supporting information is available free of charge.

MSD vs lag-times plot for 2CTD-non-phos; Cluster distribution and cluster size percentages in 2CTD crowded models at 17 mM and 25 mM protein concentrations at different salt concentrations; average R_g_ density distributions for 2CTD crowded models at 17 mM and 25 mM protein concentrations at different salt concentrations; free energy landscapes from PCA and sets of low energy conformations for 2CTD-non-phos, 2CTD-2P-5P-9P-12P, and 2CTD-2P-5P at 8.3 mM protein concentration and different salt concentrations; distributions of the number of total, intra-peptide; inter-peptide H-bonds at 8.3 mM protein concentration at different salt concentrations; contact map for 2CTD-non-phos at 8.3 mM protein concentration and 150 mM NaCl concentration; self-diffusion coefficients of 17 mM and 25 mM 2CTDs at different salt concentrations.

## AUTHOR INFORMATION

### Notes

The authors declare no competing financial interest.

## ACKNOWLEDGMENTS

The authors used the computational resources at the MERCED and PINNACLES clusters at the University of California Merced and the Advanced Cyberinfrastructure Coordination Ecosystem: Services & Support (ACCESS) under the grant TG-BIO210145. The seed grant from the UC Merced Committee on Research supported this work.

## For Table of Contents Only

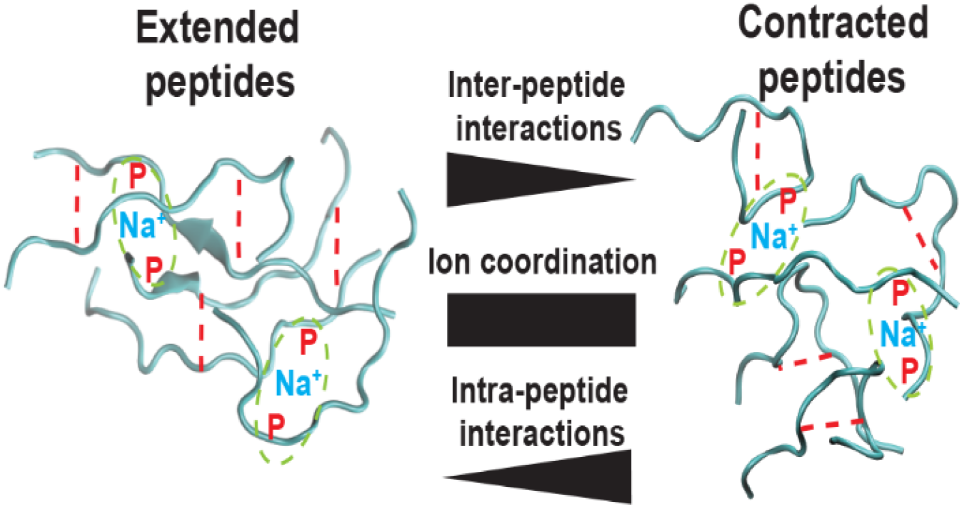

