## Supporting Information for "Clustering of RNA Polymerase II C-Terminal Domain Models upon Phosphorylation"

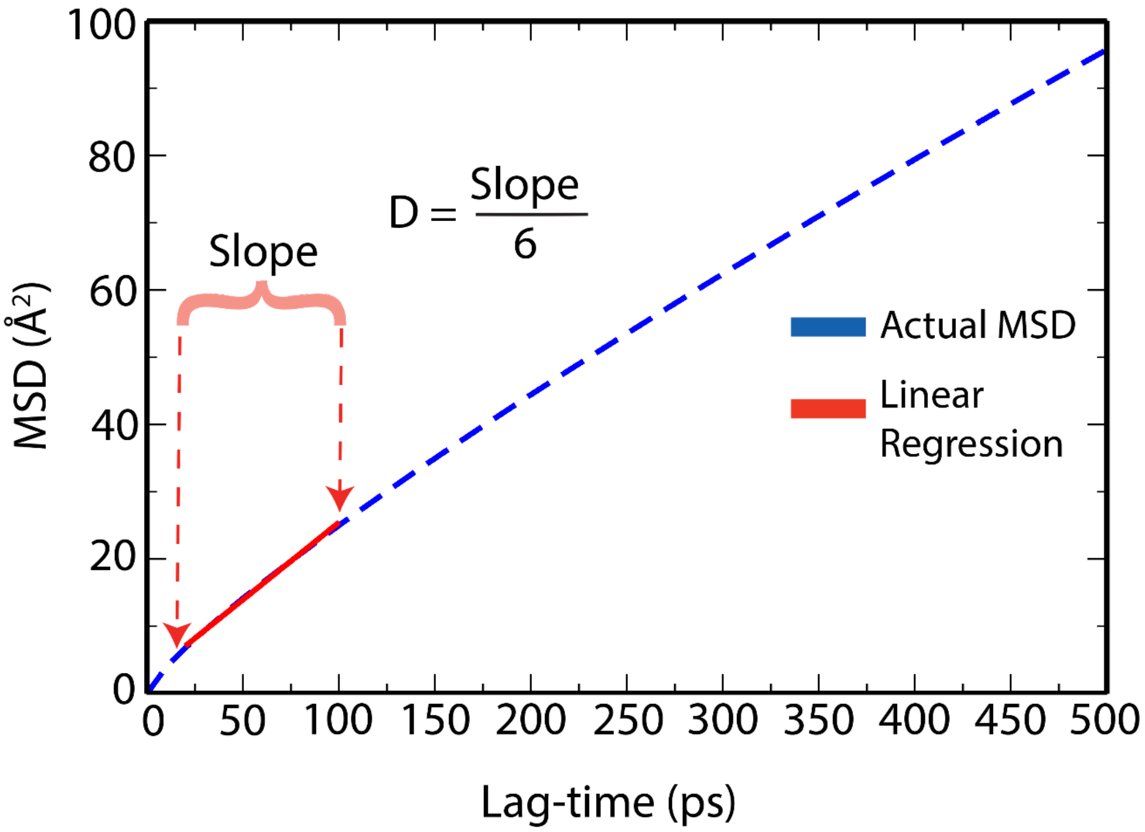


**Figure S1:** Mean-squared displacement (MSD) vs the lag-time utilized to determine slope to calculate the self-diffusion coefficient (D) for 2CTD-non-phos at 8.3 mM protein and 150 mM NaCl concentrations. Actual MSD values are shown in a blue dashed line, and the solid red line represents the fitted values to the actual MSD values with linear regression between 20 ps and 100 ps with 10 ps time intervals. The same protocol was followed to determine the self-diffusion coefficients for other 2CTD systems in Table 2, Table S1 and Table S2.

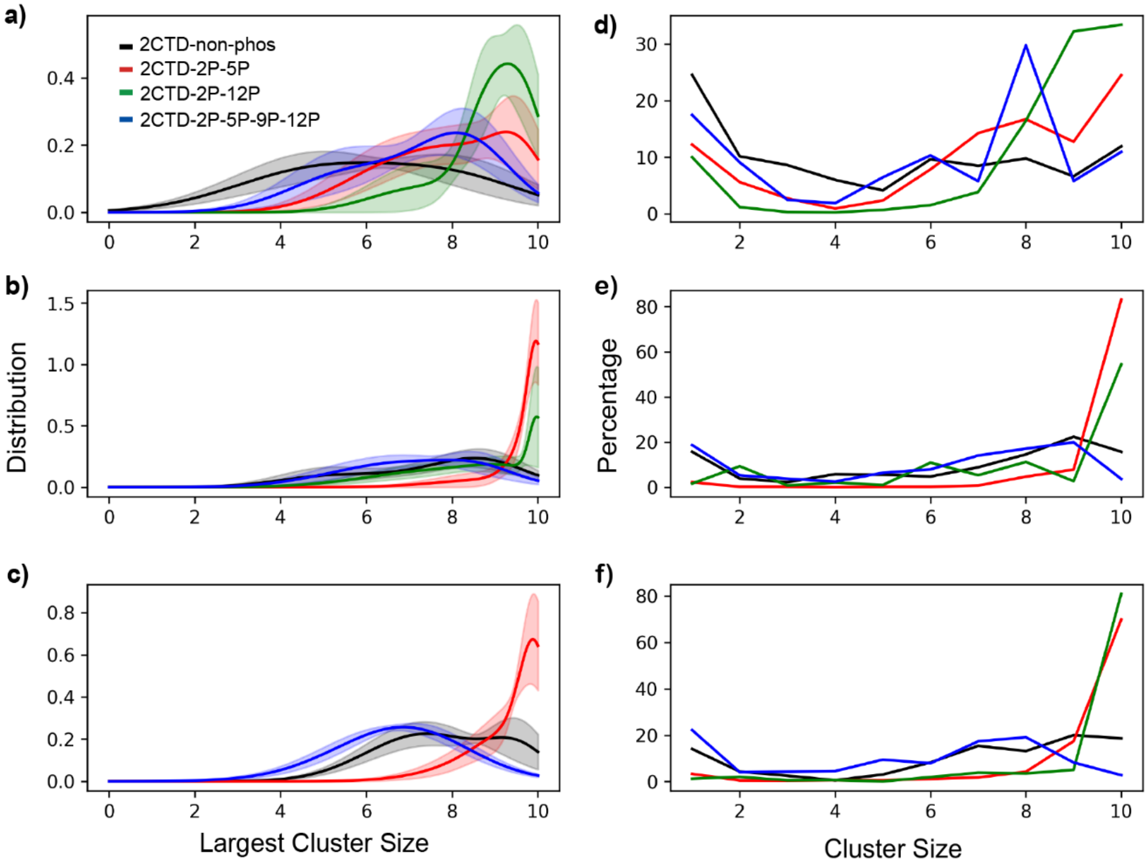


**Figure S2:** Cluster distribution and cluster size percentages in 2CTD crowded systems at 17 mM protein concentration. The largest cluster size distributions are shown at a) 0, b) 150, and c) 300 mM NaCl. Percentages of each cluster size are shown at d) 0, e) 150, and f) 300 mM NaCl. Standard errors [for panels (a), (b), and (c)] were calculated by splitting the last 800 ns of the 1 μs trajectory into 160 ns small trajectories for each system. For 2CTD-2P-12P the largest cluster size distribution is not shown in panel (c), because for most of the frames, the largest cluster size remained at 10 (also see panel (f)). The colors of the curves are the same as in panel (a) for all the other panels.

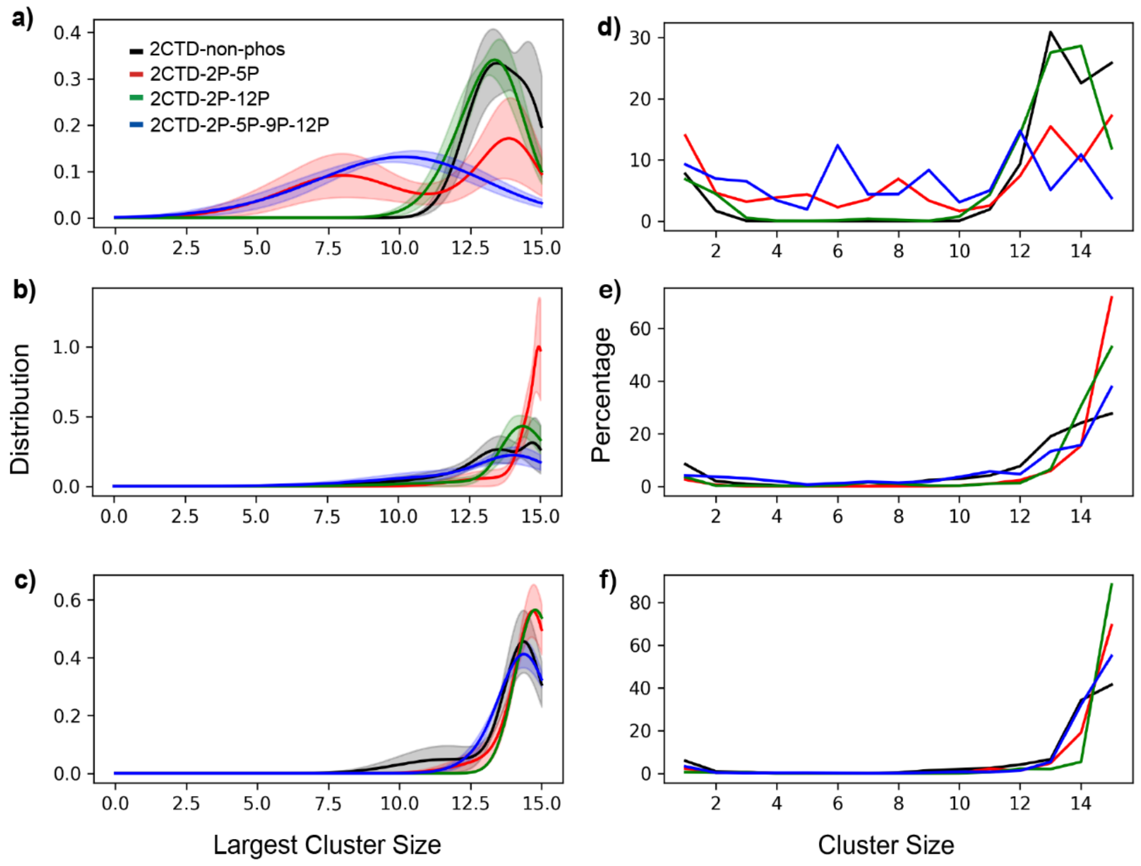


**Figure S3:** Cluster distribution and cluster size percentages in 2CTD crowded systems at 25 mM protein concentration. The largest cluster size distributions are shown at a) 0, b) 150, and c) 300 mM NaCl. Percentages of each cluster size are shown at d) 0, e) 150, and f) 300 mM NaCl. Standard errors [for panels (a), (b), and (c)] were calculated by splitting the last 800 ns of the 1 μs trajectory into 160 ns small trajectories for each system. For 2CTD-2P-12P in panel (c) the standard error is not shown because approximately between 320 ns and 480 ns of the last 800 ns the largest cluster size remained constant at 15. The colors of the curves are the same as in panel (a) for all the other panels.

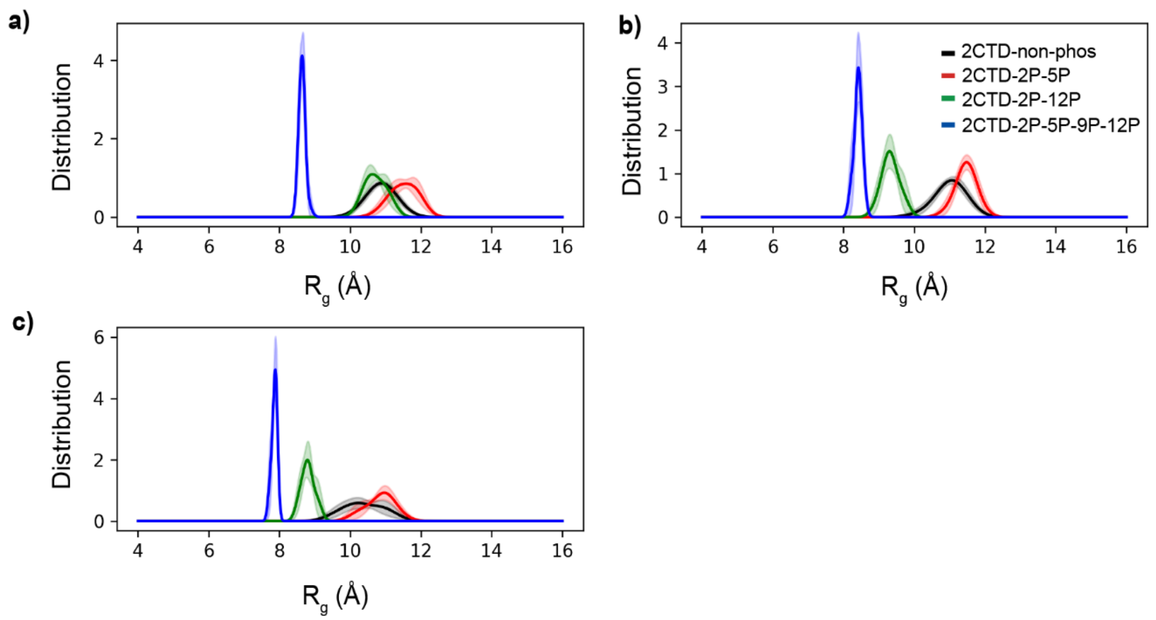


**Figure S4.** Average radius of gyration ($R_g$) density distributions for 2CTD crowded models at 17 mM protein and (a) 0, (b) 150, and (c) 300 mM NaCl concentrations. The colors of the curves are the same as in panel (b) for all the other panels. Standard errors were calculated by splitting the last 800 ns of the 1 μs trajectory into 160 ns small trajectories for each system.

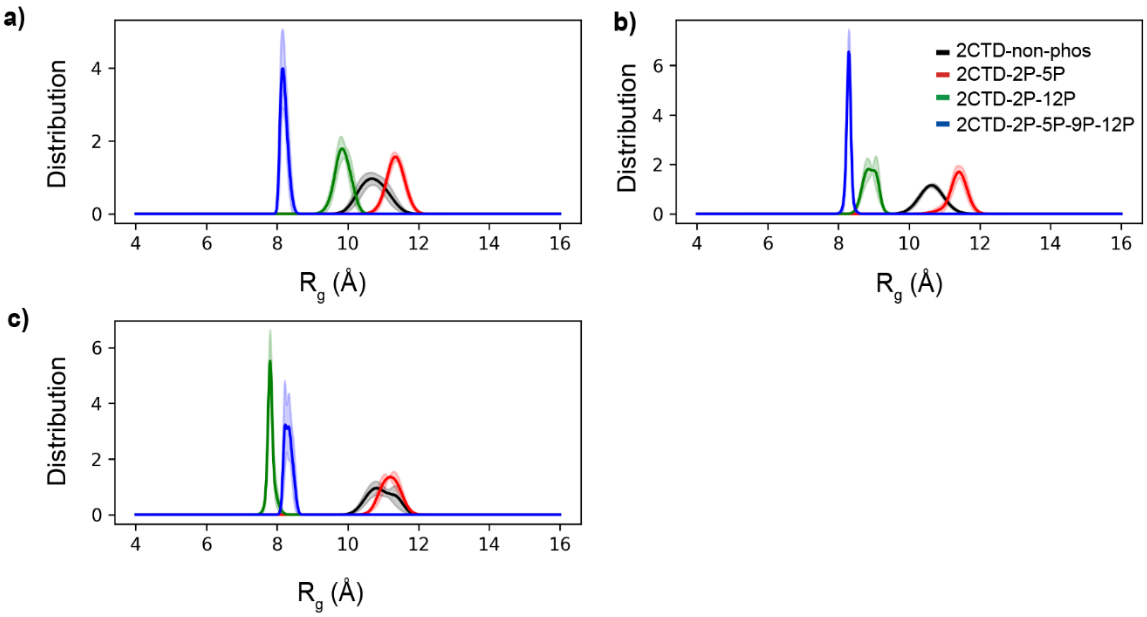


**Figure S5.** Average radius of gyration ($R_g$) density distributions for 2CTD crowded models at 25 mM protein and (a) 0, (b) 150, and (c) 300 mM NaCl concentrations. The colors of the curves are the same as in panel (b) for all the other panels. Standard errors were calculated by splitting the last 800 ns of the 1 μs trajectory into 160 ns small trajectories for each system.

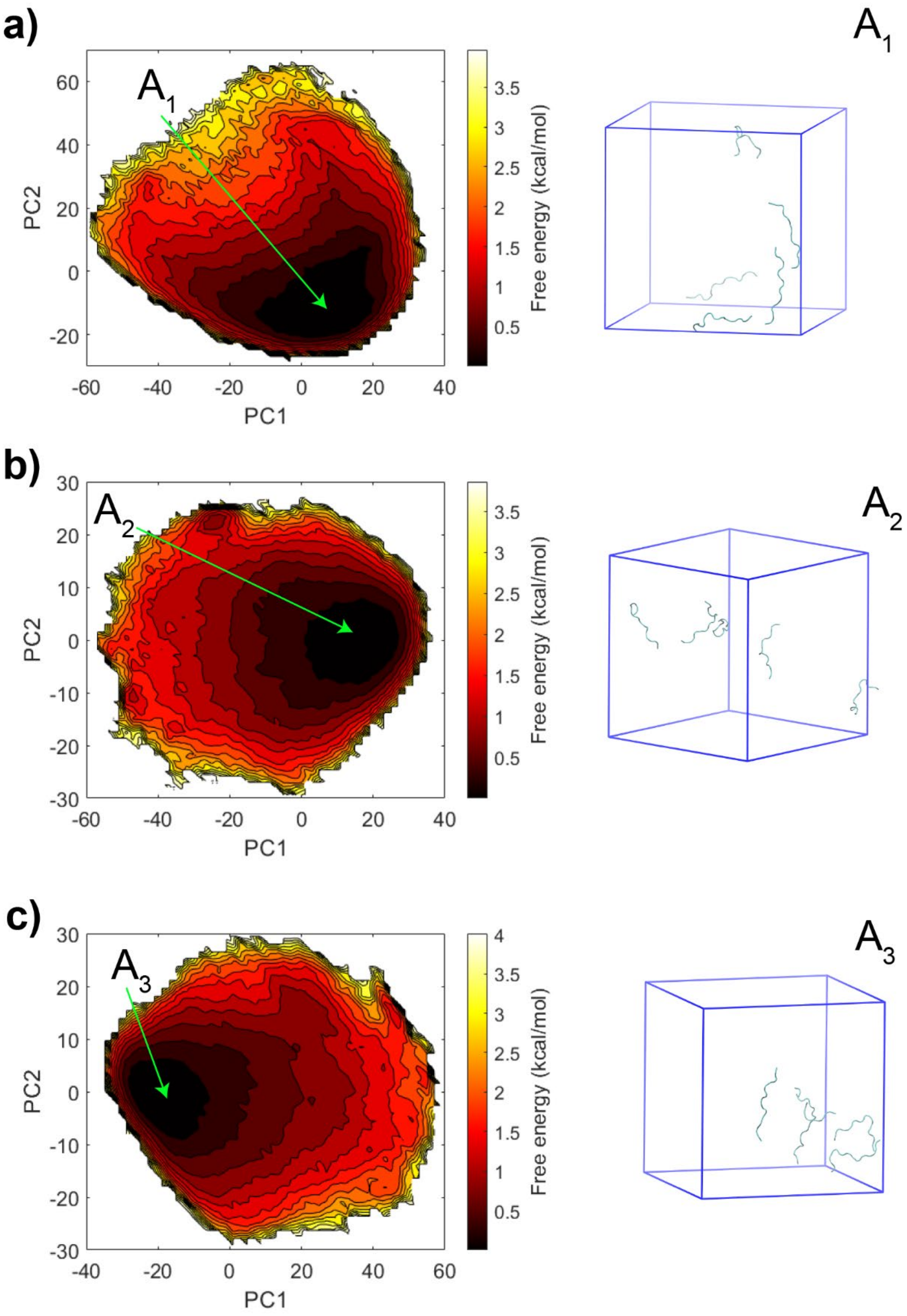


**Figure S6.** Free energy landscapes using PC1 and PC2 as reaction coordinates from the PCA using cartesian coordinates for 2CTD-non-phos crowded models at 8.3 mM protein and (a) 0, (b) 150, and (c) 300 mM NaCl concentrations. In addition, $A_1$-$A_3$ represent the frames that have low energy conformations at each salt concentration.

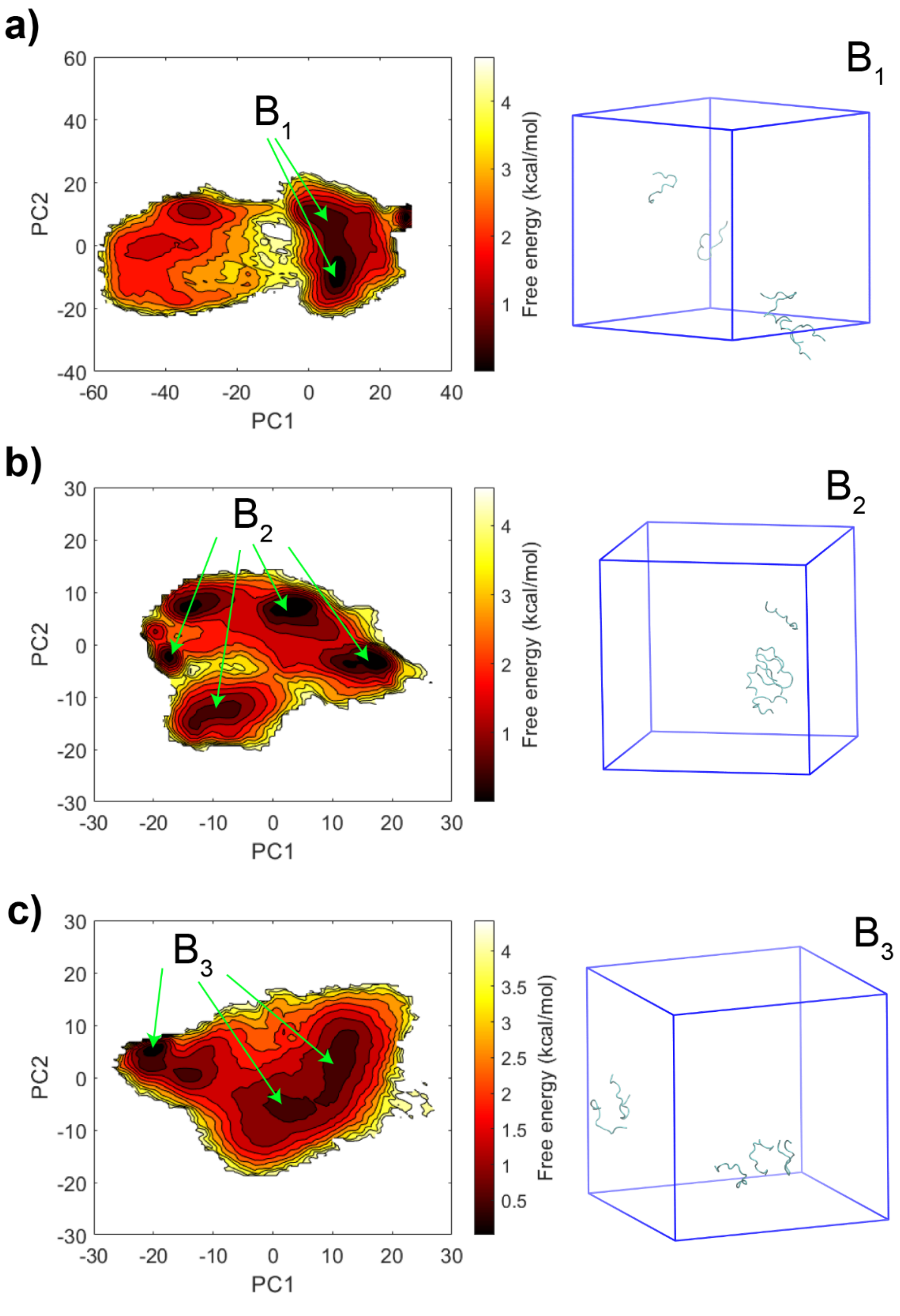


**Figure S7.** Free energy landscapes using PC1 and PC2 as reaction coordinates from the PCA using cartesian coordinates for 2CTD-2P-5P-9P-12P crowded models at 8.3 mM protein and (a) 0, (b) 150, and (c) 300 mM NaCl concentrations. In addition, $B_1$-$B_3$ represent the frames that have low energy conformations at each salt concentration. $B_1$ has conformations at two energy minima, $B_2$ has conformations at four energy minima and $B_3$ has conformations at three energy minima.

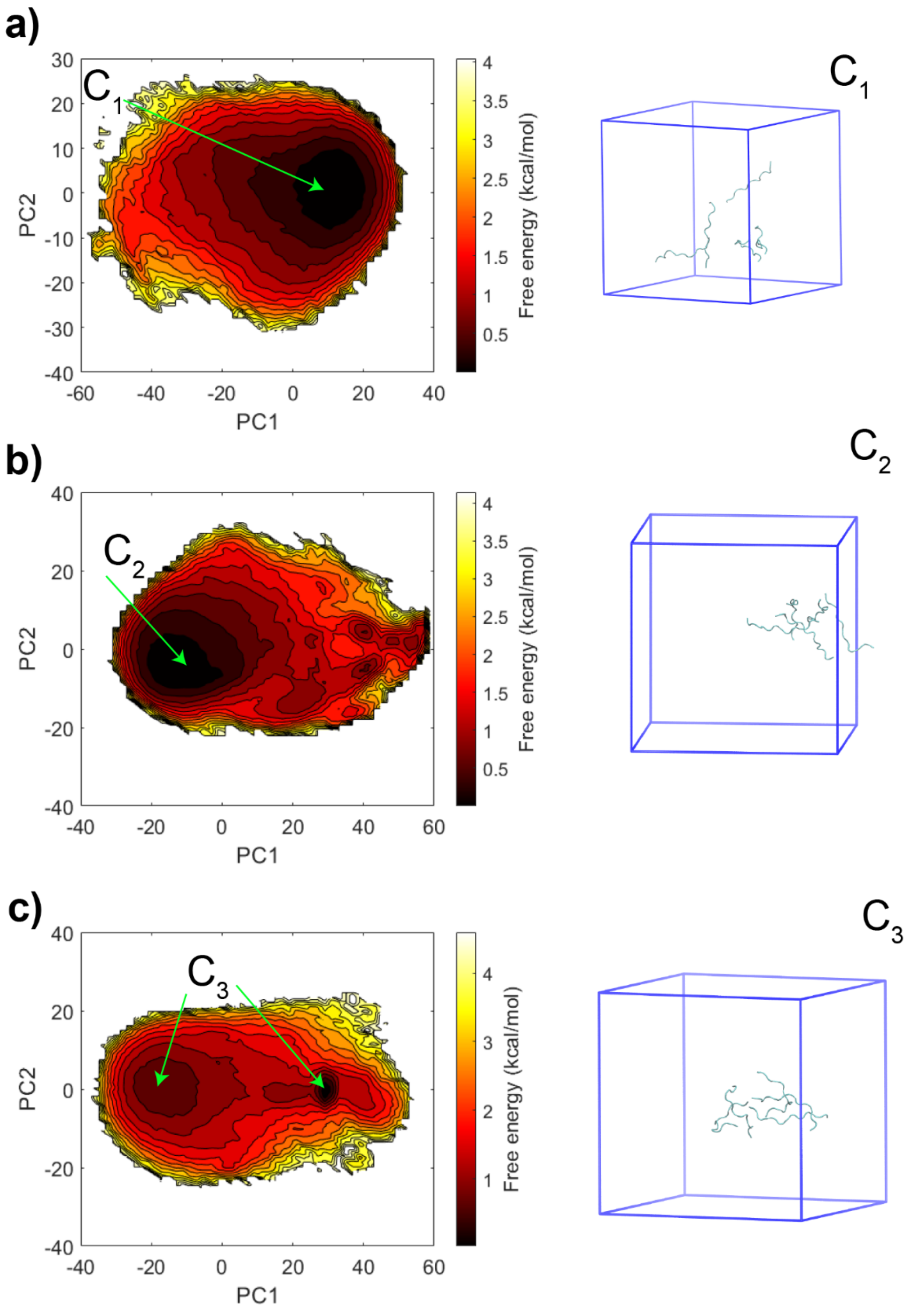


**Figure S8.** Free energy landscapes using PC1 and PC2 as reaction coordinates from the PCA using cartesian coordinates for 2CTD-2P-5P crowded models at 8.3 mM protein and (a) 0, (b) 150, and (c) 300 mM NaCl concentrations. In addition, $C_1$-$C_3$ represent the frames that have low energy conformations at each salt concentration. $C_3$ has conformations at two energy minima.

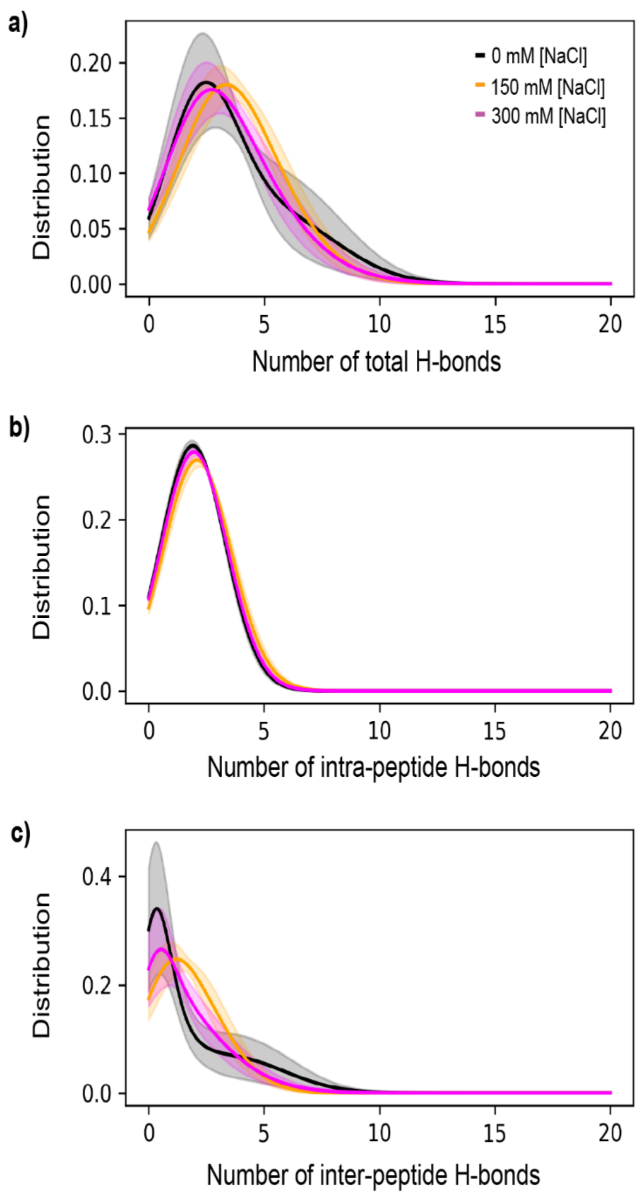


**Figure S9.** Distributions of the number of (a) total, (b) intra-peptide, and (c) inter-peptide H-bonds for 2CTD-non-phos system at 8.3 mM protein and different salt concentrations. The colors of the curves are the same as in panel (a) for all the other panels. Standard errors were calculated by splitting the last 800 ns of the 1 μs trajectory into 160 ns small trajectories for each system.

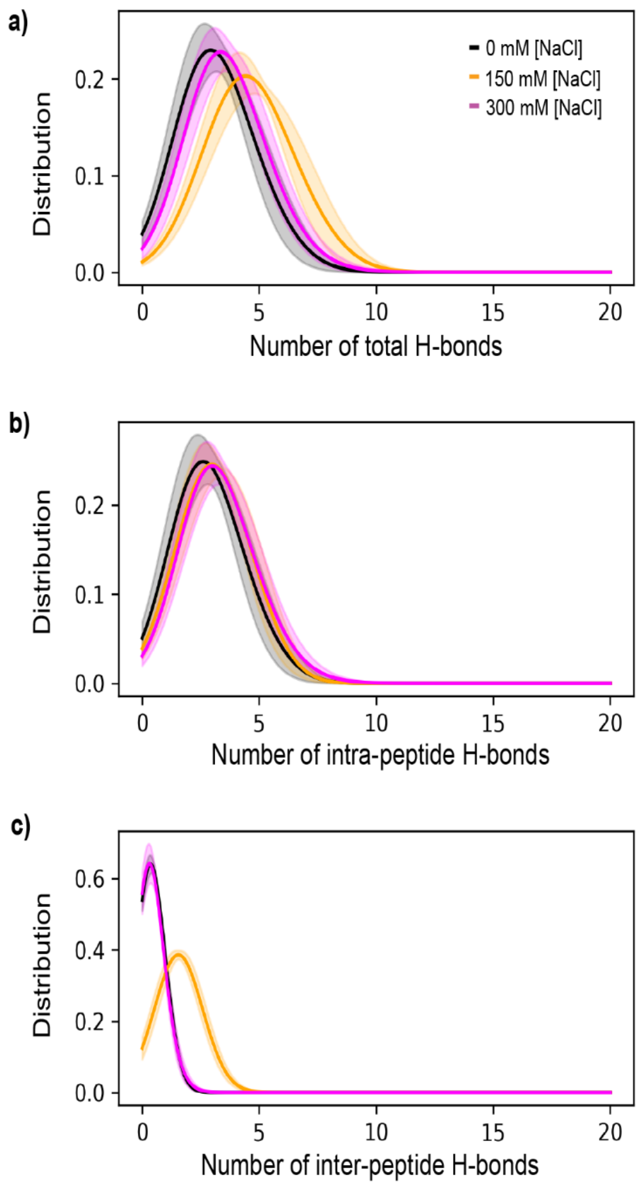


**Figure S10.** Distributions of the number of (a) total, (b) intra-peptide, and (c) inter-peptide H-bonds for 2CTD-2P-5P-9P-12P systems at 8.3 mM protein and different salt concentrations. The colors of the curves are the same as in panel (a) for all the other panels. Standard errors were calculated by splitting the last 800 ns of the 1 μs trajectory into 160 ns small trajectories for each system.

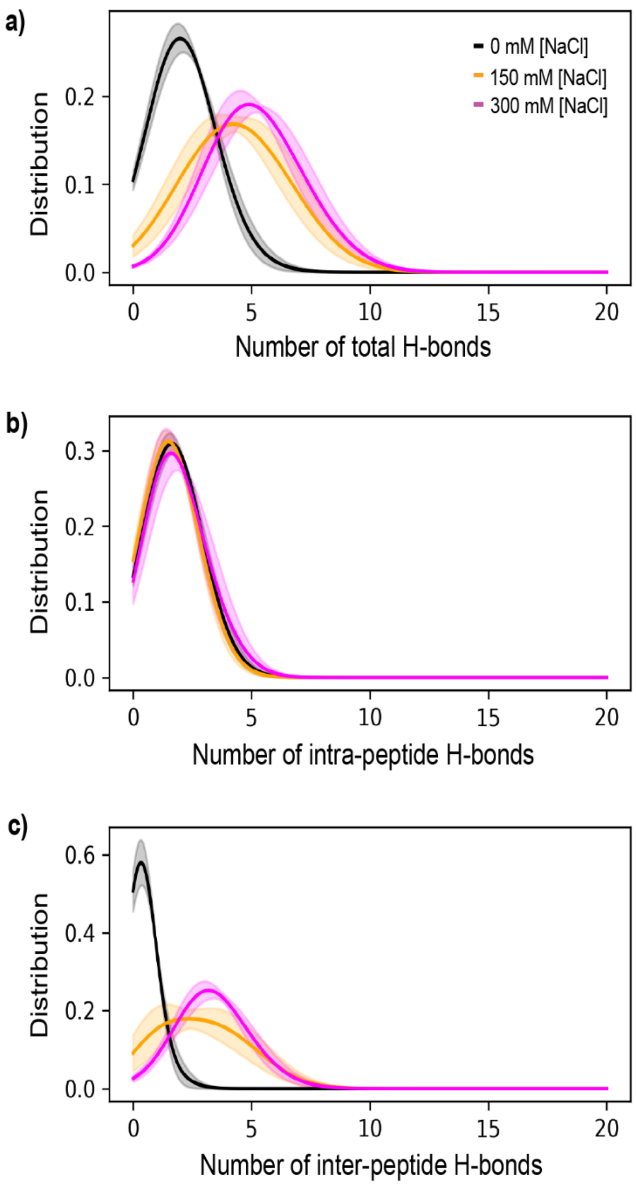


**Figure S11.** Distributions of the number of (a) total, (b) intra-peptide, and (c) inter-peptide H-bonds for 2CTD-2P-5P systems at 8.3 mM protein and different salt concentrations. The colors of the curves are the same as in panel (a) for all the other panels. Standard errors were calculated by splitting the last 800 ns of the 1 μs trajectory into 160 ns small trajectories for each system.

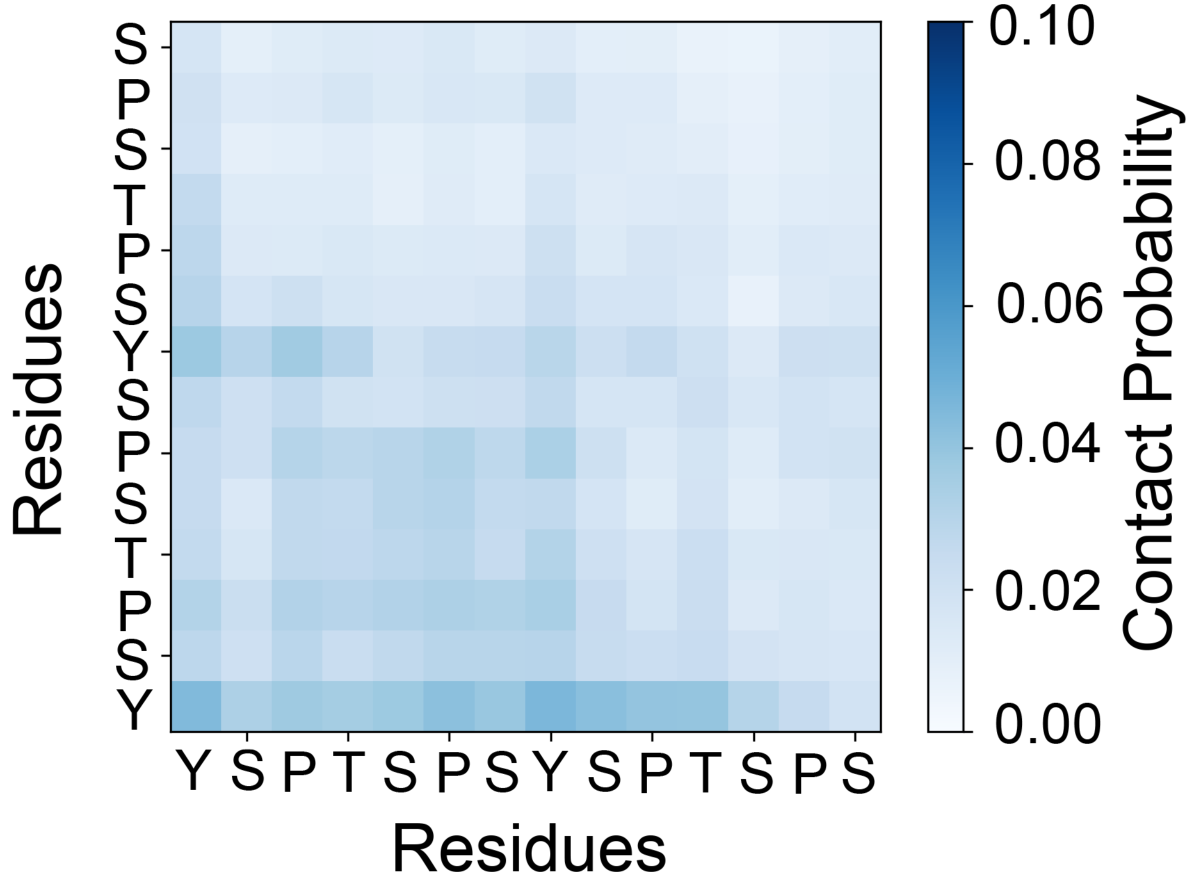


**Figure S12.** Contact map for inter-peptide contacts between residues within 5 Å for the 2CTD-non-phos system at 8.3 mM protein and 150 mM NaCl concentrations. Contacts were averaged for each residue pair over the trajectory and the number of peptide pairs.

**Table S1.** Self-diffusion coefficients of 17 mM 2CTDs at different salt concentrations.

| NaCl concentration (mM) | Diffusion Coefficients ($10^{-2}$ Å$^2$ ps$^{-1}$) | | | |
|---|---|---|---|---|
| | 2CTD-non-phos | 2CTD-2P-5P | 2CTD-2P-12P | 2CTD-2P-5P-9P-12P |
| 0 | 3.05 | 2.0 | 1.45 | 1.53 |
| 150 | 2.54 | 1.38 | 1.31 | 1.56 |
| 300 | 2.28 | 1.37 | 1.07 | 1.50 |

**Table S2.** Self-diffusion coefficients of 25 mM 2CTDs at different salt concentrations.

| NaCl concentration (mM) | Diffusion Coefficients ($10^{-2}$ Å$^2$ ps$^{-1}$) | | | |
|---|---|---|---|---|
| | 2CTD-non-phos | 2CTD-2P-5P | 2CTD-2P-12P | 2CTD-2P-5P-9P-12P |
| 0 | 1.71 | 1.83 | 1.28 | 1.28 |
| 150 | 1.82 | 1.17 | 0.99 | 0.96 |
| 300 | 1.62 | 1.14 | 0.76 | 0.74 |
